# Triglyceride/Cholesterol Ester Ratio Encodes Lipid Droplet Size and Diversity

**DOI:** 10.64898/2026.01.21.700800

**Authors:** Helin Elhan, Calvin Dumesnil, Mehdi Zouiouich, Cyril Moulin, Alenka Čopič, Mohyeddine Omrane, Bleuenn Feuillet, Abdou Rachid Thiam

## Abstract

Lipid droplet (LD) heterogeneity is a hallmark of their pathophysiological relevance. This is especially evident when cells sequester toxic cholesterol by converting it into cholesterol esters (CEs) within LDs. Since CEs can form liquid crystals (LCs), it remains unclear how such ordered structures are accommodated within the inherently dynamic environment of LDs. Here, we show that the fluidizing properties of triglycerides (TGs) help CE incorporation and influence LD growth and heterogeneity. Seipin, the key regulator of TG-LD formation and size, does not significantly impact CE-rich LD size. Instead, the CE/TG ratio, the sequence of neutral lipid deposition, and the activity of diacylglycerol acyltransferases, especially DGAT2, determine whether LDs enlarge, remain fluid, or lock into LC phases. We found that the LC phase resists LD ripening and acts as a kinetic barrier to lipid entry. Lastly, we observe that perilipins, the most abundant LD surface proteins, differentially target CE-rich LDs: Plin3 and 5 are excluded from them, whereas Plin2 and 4 are favored. These findings highlight the CE/TG ratio as a key organizing principle of lipid storage and LD function, with immediate relevance for diseases linked to sterol accumulation.

## Introduction

Cholesterol homeostasis is vital for cellular function and overall health, especially in the liver and brain, which are key organs for cholesterol metabolism. While cholesterol is necessary for maintaining membrane integrity and producing bile acids, excess free cholesterol can be toxic to cells. To prevent this, cells convert cholesterol into cholesterol esters (CE), which are stored in lipid droplets (LDs). This process also provides a flexible, adaptive way to manage changes in cholesterol levels. However, an imbalance in cholesterol intake, esterification, and removal can cause harmful buildup of sterols and CEs, contributing to metabolic disorders such as hypercholesterolemia and metabolic syndrome. Furthermore, in Alzheimer’s disease (AD), CE accumulation occurs early, including within neuronal LDs. In iPSC-derived AD neurons, higher CE levels promote the buildup of phosphorylated tau, a key disease marker, and enhance alpha-synuclein’s association with LDs. Blocking CE synthesis by genetic manipulation of acetyl-CoA acetyltransferase 1, which converts free cholesterol into CE, significantly reduces tau pathology in the AD mouse model. Despite the critical role of CE in disease, the molecular mechanisms regulating CE storage and its phase behavior within LDs remain poorly understood.

Improper sequestration of cholesterol into CE-rich LDs can form cholesterol crystals^7^, which trigger inflammation and contribute to conditions such as non-alcoholic steatohepatitis and atherosclerosis ^1,8^. Notably, CE can form liquid-crystalline phases within LDs and may crystallize in the presence of solid nuclei^9^, a phenomenon similar to that observed in cells from atherosclerotic patients^7^. There, CE-rich LDs are connected to crystal fibers likely containing CE and cholesterol. Therefore, understanding the mechanisms governing CE LD composition and organization is crucial for comprehending cholesterol regulation and its impact on the development of metabolic and neurodegenerative diseases.

LDs are widely present across organisms^10–12^ and play key roles in energy storage, lipid metabolism, and cellular homeostasis^12,13^. They are highly dynamic organelles that enable cells to efficiently store and mobilize lipids in response to metabolic demands and various types of stress^14,15^. LDs form within the endoplasmic reticulum (ER) when hydrophobic neutral lipids, mainly triglycerides (TGs), are fabricated and eventually undergo phase separation within the bilayer^11,16–18^. This nucleation step is controlled by a key protein called seipin, which forms a donut-shaped oligomer wherein the nascent LD initiates. Not only does seipin nucleate TG LDs, but it also ensures their proper growth by preventing, at least in part, the loss of TG during ripening^16,17^. During ripening, small LDs with higher internal Laplace pressure release their TG content to the bilayer, which is then taken up by larger LDs with lower pressure. LDs can nucleate without seipin, albeit erratically, and abnormally large LDs appear. By controlling nucleation and growth, seipin enables the formation of relatively uniform-sized TG LDs.

The story becomes complex when cells produce CE. We recently demonstrated that TGs, which interact favorably with seipin, create a fluid environment that favors the nucleation of CE LDs at seipin sites^9^. Mechanisms of CE LD nucleation that rely on TG remain elusive. Also, whether seipin regulates CE LD growth and function as it does for TG LDs is unknown.

TGs and CEs exhibit significant physicochemical differences. For instance, triolein melts at around 4°C and cholesterol oleate at approximately 44°C^9,19^. The distinct chemical properties of TGs and CEs significantly influence LD behavior^19^. Recent insights have revealed that, at specific CE concentrations in LDs, the LD core can become metastable and eventually lock into a liquid-crystalline (LC) phase^9,20–22^. These results suggest that smaller CE droplets offer greater thermodynamic stability. This shift affects the lipid core matrix and may also modulate the binding of LD-associated proteins, underscoring the dynamic interplay between lipids and proteins in shaping LD function^21,23–27^.

TGs melt CE as salt melts ice, highlighting the relevance of regulating the CE/TG ratio in cells. This melting effect is nicely illustrated in lipoproteins, with increasing CE levels as lipoprotein size decreases^28^. CE/TG ratio mirrors the lipoprotein structure, function, and metabolic fate. This ratio influences not only the physical properties of lipoproteins, such as size, density, and surface curvature, but also their biological roles in lipid transport, energy delivery, and cholesterol homeostasis, which are determined by the different apolipoproteins that surround them. For instance, TG-rich lipoproteins like chylomicrons and very low-density lipoproteins (VLDL) primarily function in energy distribution, delivering lipids to peripheral tissues^28,29^. In contrast, CE-rich lipoproteins, such as low-density lipoproteins (LDL) and high-density lipoproteins (HDL), are critical for cholesterol transport to and from tissues, respectively ^28,29^. Alterations in the CE/TG ratio have been implicated in dyslipidemias, atherosclerosis, and metabolic diseases^30–32^, underscoring its importance as both a functional parameter and a potential biomarker of lipid metabolism.

The CE/TG ratio can vary significantly within LDs^9,22,33^, and while its relevance to lipoprotein metabolism and pathophysiological conditions is relatively well established, its contribution to LD biology remains largely unclear. Adipocytes predominantly store TGs^34^; macrophages can exhibit highly CE-enriched LDs^35,36^. Notably, this diversity extends beyond cell type. Even within a single cell, TG-rich and CE-rich LD subpopulations may coexist, as in steroidogenic cells^23^, and be spatially distinct, with CE LDs potentially forming closer to the nuclear envelope in A431 cells^37^. This intracellular LD heterogeneity may affect protein association with LDs, notably with perilipins, the major LD proteins, suggesting distinct functional roles for TG- and CE-rich LDs, which remain poorly understood.

Our previous work highlighted an overlooked function of TG beyond its energy and lipid storage functions^9^. It acts as a nucleator of CE LD at seipin. This prompted us to investigate how seipin and TGs further control the incorporation of CE into LDs and how LD composition affects the recruitment of the most abundant LD proteins in a model cellular system.

## Results

### Mature CE LD size and crystallinity are mainly determined by triglycerides rather than seipin

We previously characterized the internal LD core state based on TG/CE ratios and defined nutrient conditions to modulate the liquid state within LDs. At 37°C, LDs exhibit three states: with low CE levels (∼20 mol%), they show an isotropic stable liquid phase; above ∼20 mol%, they become metastable but can exhibit an isotropic phase; above ∼90 mol%, they are in a Liquid Crystalline phase (LC) (**Fig. 1A**). Under polarized light, CE-rich LDs show a characteristic Maltese cross motif^9^.

**Figure 1.**
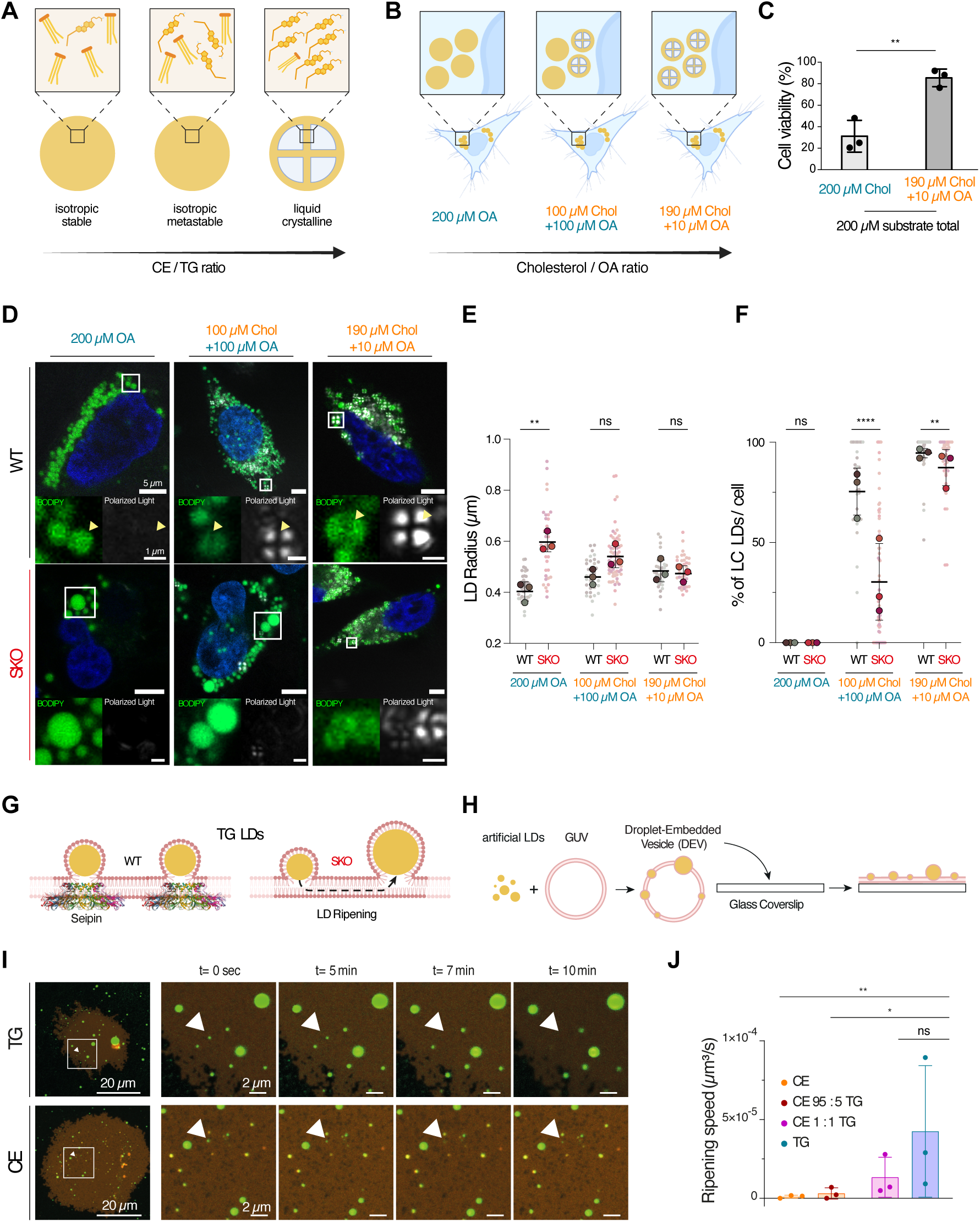
Lipid composition influences LD phase behavior, seipin-mediated size regulation, and resistance to ripening. **(A)** Nutrient composition affects the liquid state of lipid droplets (LDs) and cell viability. A schematic illustrates the LD state as a function of the CE/TG ratio at 37°C. At low CE/TG ratios (≈20 mol%, left), LDs are rich in TG and remain isotropic and stable. At intermediate CE/TG ratios (up to ≈90 mol%, center), LDs become metastable and CE-rich but still isotropic. At higher CE/TG ratios (>90 mol%, right), LDs transition to a liquid-crystalline (LC) phase, marked by the appearance of a distinct Maltese cross pattern under polarized light. **(B)** Schematic showing LD states in cells as a function of the Cholesterol/Oleic Acid (Chol/OA) ratio in the surrounding environment. Loading with 200 µM OA leads to the formation of TG-rich LDs without an LC phase (left). Co-loading with 100 µM Chol and 100 µM OA results in a mixed population of LDs, including both LC-LDs and TG/CE isotropic LDs (center). Loading with 190 µM Chol and 10 µM OA produces CE-rich LDs with a characteristic LC phase (right). **(C)** The percentage of HeLa wild-type (WT) cell viability after 24 hours of lipid loading with 200 µM total substrate, using either 200 µM Chol alone or 190 µM Chol + 10 µM OA. The presence of 10 µM OA significantly reduces cholesterol-induced cell death. Each data point shows the mean of an independent experiment (*N* = 3). Bars represent mean ± SD. Statistical significance was assessed with Student’s *t*-test: \*\**p* = 0.0051. **(D)** Seipin’s effect on LD size and phase varies between TG and CE LC LDs. Representative images of HeLa WT and Seipin knockout (SKO) cells after 24 hours of lipid loading with OA or combined OA and Cholesterol at the specified concentrations. Polarized light and BODIPY staining reveal distinct LD subpopulations with either isotropic or LC internal organization. Nuclei are stained with Hoechst. Yellow arrows indicate the presence or absence of polarized light signals. Each experiment was repeated three times with similar results. **(E)** Average LD size per cell was quantified. Data are shown as SuperPlots, with small dots representing individual cell means and large dots representing the mean of each independent replicate (*N* = 3). Bars indicate mean ± SD. Significance was tested by Student’s *t*-test: 200 µM OA: WT *n* = 40, SKO *n* = 37, \*\**p* = 0.0033; 100 µM Chol + 100 µM OA: WT *n* = 34, SKO *n* = 66, not significant (ns), *p* = 0.0589; 190 µM Chol + 10 µM OA: WT *n* = 35, SKO *n* = 38, ns, *p* = 0.7429. **(F)** Fraction of LC LDs per cell was analyzed and shown as SuperPlots, with small dots for individual cell means and large dots for the mean of each replicate (*N* = 3). Bars display mean ± SD. Significance was determined with Mann–Whitney test: for 200 µM OA: WT *n* = 34, SKO *n* = 38, ns, *p* = NA; 100 µM OA + 100 µM Chol: WT *n* = 34, SKO *n* = 66; 190 µM Chol + 10 µM OA: WT *n* = 35, SKO *n* = 38, \*\**p* = 0.0013. **(G)** CE-rich artificial lipid droplets are less prone to ripening. Diagram of seipin’s role during TG LDs growth in the ER. Left: In wild-type cells, seipin (shown in side view) binds at the bottom-neck of forming LDs, aiding proper TG packaging. Right: In seipin knockout cells, TG LDs tend to ripen, leading to destabilization that causes smaller LDs to shrink and larger ones to grow bigger. **(H)** Schematic of the droplet-embedded vesicles (DEVs) protocol. Artificial LDs (aLDs), either TG or CE, are mixed with PC/PE + cy-5PE (70%/30% + 0.5%) GUVs and BODIPY. The aLDs insert into the leaflets of the GUV bilayer, forming a DEV. BODIPY stains the droplets. These DEVs are then placed on an untreated glass coverslip where they rupture and flatten into a 2D shape. **(I)** Time course of ripening in flattened DEVs containing aLDs made of pure TG or CE. **(J)** Ripening rate of flattened DEVs was determined by tracking droplet size over 1 hour and fitting the data linearly. Each point reflects the mean of one experiment. Data are shown as mean ± SEM. CE-rich aLDs (95 mol% CE / 5 mol% TG) ripen 5–15 times slower than equimolar TG/CE or pure TG droplets (\*\**p* = 0.0077, \**p* = 0.0359, ns *p* = 0.3642).

Loading HeLa cells with 200 µM Oleic Acid (OA) for 24 hours triggered TG-rich LDs (**Fig. 1B**). Supplying the same amount of cholesterol via methyl-β-cyclodextrin (MβCD) caused significant cell death from cholesterol toxicity (**Fig. 1C**). Remaining cells’ LDs developed an LC phase. Remarkably, adding 10 µM OA to 190 µM cholesterol significantly improved cell viability and promoted LC LD formation (**Fig. 1B-C**). This small OA dose accelerates nucleation of CE-rich LDs, likely via TG synthesis^9^. These data suggest the initial TG/CE ratio during CE storage is critical to cell fate when cholesterol becomes in excess. When cells were supplemented with equal amounts of OA and cholesterol (100 µM each), mixed LDs comprising LC and isotropic TG/CE LDs were produced (**Fig. 1B**).

Building on these settings, we investigated how the TG/CE ratio affects LD growth and how seipin regulates this process. We used HeLa WT and seipin KO (SKO) cells in three different nutrient-feeding conditions for 24 hours: 200 µM of OA (hereafter referred to as OA), 10 µM of OA and 190 µM of cholesterol (referred to as cholesterol), and 100 µM of OA and 100 µM of cholesterol (referred to as equimolar) (**Fig. 1D**). Imaging was conducted at 37°C.

For clarity, we use the following terminology: TG LDs for LDs made in OA, most of which are in an isotropic and stable liquid state; CE LDs for LDs made with cholesterol, most of which exhibit LC organization; non-LC and LC LDs made in equimolar concentration, where LC LDs are probably highly enriched in CE and the non-LC are more enriched in TG.

In OA feeding, WT cells exhibited homogeneous TG LDs (**Fig. 1D, Supplementary Fig. 1A**). The absence of seipin significantly affected TG LD size distribution, leading to larger LDs (**Fig. 1E**). Under cholesterol-fed conditions, nearly all LDs had an LC structure in both WT and SKO cells (**Fig. 1F**). Surprisingly, seipin KO showed no impact on LD size distribution, with similar sizes in WT and KO cells (**Fig. 1E**), suggesting the protein does not impact CE LD size as it does for TG LDs.

Using BODIPY dye, differences in staining were observed between TG LDs and CE LDs (**Fig. 1D**). BODIPY uniformly stained TG LDs, while CE LDs showed a bracket-like pattern. This signal intensity difference may result from BODIPY’s partial exclusion from the LC lattice of CE LDs, or changes in quantum efficiency or diffraction patterns due to the LC environment. Alongside polarized light imaging, the BODIPY signal pattern could also serve as a proxy for the internal structure of LDs.

The equimolar OA/cholesterol condition exhibited more complex phenotypes. Non-LC and LC LDs coexisted within a single cell, suggesting different TG/CE loads among LD subpopulations. Initially, the data showed minimal differences in size distribution between WT and SKO cells (**Fig. 1E, Supplementary Fig. 1B**), although the frequency of larger LDs increased, sometimes exceeding the size of TG LDs. Further analysis revealed subtle effects of seipin absence. The size difference was significant between WT and SKO in the non-LC LDs, similar to the difference observed in TG LDs (**Supplementary Fig. 1C**). LC LDs did not display substantial differences, paralleling the case of CE LDs induced by cholesterol feeding. Therefore, within individual cells, two subpopulations coexisted during equimolar feeding: non-LC LDs, whose size depended on seipin presence (resembling pure TG LDs), and LC LDs, whose size was independent of seipin (similar to the pure CE LD case).

Analyzing the proportion of LC LDs within the LD subpopulation under this equimolar condition revealed a 3-fold decrease in the proportion of LC LDs in SKO cells compared to the WT condition (**Fig. 1F**). This suggests that the absence of seipin increases the TG/CE ratio in LDs, decreasing the fraction of LC LDs. Additionally, we occasionally observed LDs displaying a hybrid or partially polarized light pattern, particularly in SKO cells, which we classified as LC LDs. These patterns indicate that these LDs possess a central TG-rich isotropic core surrounded by a CE-rich LC shell (**Supplementary Fig. 1D**). Consistently, a uniform Bodipy signal was detected in the core of these LDs. This suggests a potential disruption of the LC phase by TGs, possibly due to increased incorporation of CEs in the periphery of non-LC LDs or increased incorporation of TGs in the core of LC LDs.

In summary, our data suggest that seipin affects the size distribution of mature TG, but to a lesser extent than CE LDs, indicating that it does not significantly influence CE LD growth. An increased TG/CE ratio causes non-LC LDs to be under seipin regulation, while decreasing this ratio makes them less dependent on seipin. Seipin’s presence promotes more LC phases, because it samples TG in more LDs, whereas its absence results in a higher TG/CE ratio as fewer large TG-rich LDs form. We further tested these conclusions.

### Cholesterol ester is a kinetic barrier to ripening

Seipin prevents TG LDs from ripening, thereby promoting their growth ^16,17^. Removing seipin causes smaller LDs to shrink. Their material is transferred to larger ones, which develop, contributing to the bimodal size distribution of TG LDs^16^ (**Fig. 1G**). Since the LC LD size was independent of seipin and depended on TG, we hypothesized that LC LDs might be less susceptible to ripening and could be regulated by TG. To test this, we examined neutral lipid flow across model LDs in a bilayer using the Droplet-Embedded Vesicle (DEV) system^38^ (**Fig. 1H**). In this system, artificial Lipid Droplets (aLDs) are embedded between the bilayer leaflets.

We made giant unilamellar vesicles (GUVs) with dioleoylphosphatidylcholine (DOPC) and dioleoylphosphatidylethanolamine (DOPE) phospholipids in a 70/30 mol% ratio, with 0.5 mol% cy5-PE to label the membrane. Then, we mixed the GUVs with artificial LDs (aLDs) containing different neutral lipid compositions to form DEVs (**Fig. 1H**). The DEVs were deposited onto glass coverslips to observe changes in aLD size over time. Upon contact with the glass surface, the DEVs ruptured, spread, and flattened. This setup enabled us to track ripening dynamics in two dimensions, with all droplets in a single focal plane (**Fig. 1I**).

By monitoring the evolution of the average aLD size in the membrane, we can assess the ripening rate^39,40^. Remarkably, we found that pure CE aLDs exhibited minimal ripening, in stark contrast to TG aLDs (**Fig. 1J**). Even very small CE aLDs, smaller than TG aLDs that ultimately disappeared due to ripening, remained stable. These results suggest that CE does not undergo ripening with the same efficiency as TG, supporting the hypothesis that seipin is not required to prevent CE LD ripening, and that ripening does not contribute to CE LD growth, unlike in the TG case.

Interestingly, increasing the TG/CE ratio within the aLDs gradually accelerated ripening (**Fig. 1J**, **supplementary Fig. 1E**), underscoring the critical role of the TG/CE ratio in controlling LD stability and growth during ripening. TG actively promotes ripening, while CE seems to prevent it, likely due to the liquid-crystalline structures it forms. Overall, these data support our view that seipin’s specific function is to regulate TG rather than CE LD growth.

### The sequence of neutral lipid deposition influences LD composition, structure, and diversity

Our data in Fig. 1, especially with the equimolar feeding, revealed interesting observations that neutral lipids within the same cells may be packaged into LD subpopulations with different internal organizations. We observed non-LC TG/CE LDs and LC-LDs, with or without Maltese crosses, but with varying internal structures as confirmed by the Bodipy dye brightness and pattern. We therefore wondered how the sequence of incorporation of TG and CE in these LDs is crucial for determining LD structure. To test this, we sequentially treated the cells with 200 µM OA for 24 hours, then with cholesterol (190 µM + 10 µM OA), or vice versa, compared to simultaneous addition of both (**Fig. 2A**). In all conditions, the same amount of substrate was provided. In each condition, we measured the number and size of LDs, the proportion of LDs with a polarized light signal, and the intensity of that signal; this intensity is low when there is no liquid crystalline phase or when LC LDs are small.

**Figure 2.**
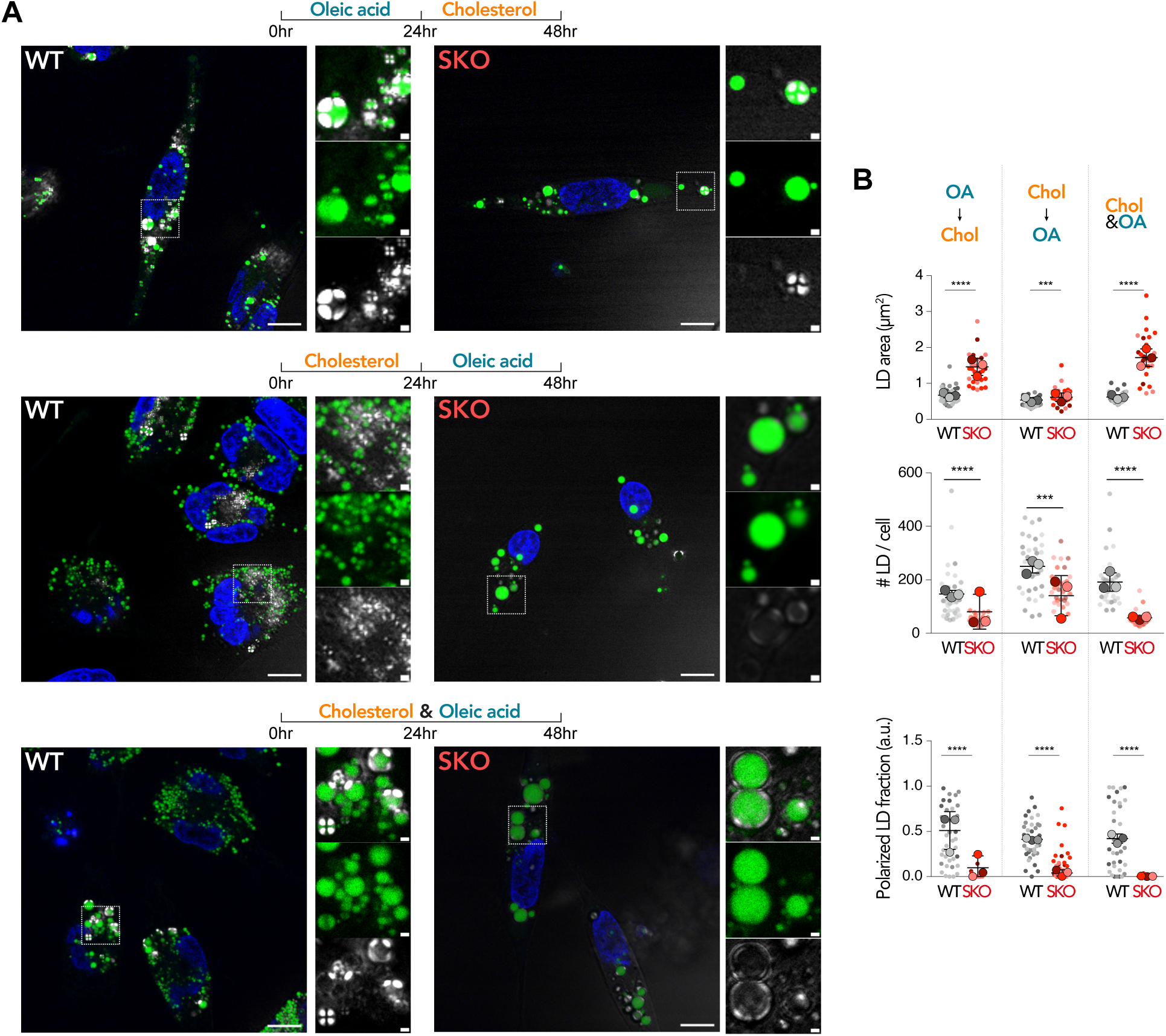
Sequential neutral lipid supply controls LD composition, internal structure, and Seipin-dependent remodeling. **(A)** Representative microscopy images of HeLa WT and Seipin knockout (SKO) cells after lipid loading under three conditions, all providing the same total substrate amount (200 µM). Cells were incubated either sequentially with 200 µM OA for 24 hours followed by 190 µM Chol + 10 µM OA for 24 hours (OA→Chol), in the reverse order (Chol→OA), or with 100 µM OA + 100 µM Chol added simultaneously (equimolar Chol&OA). Polarized light and BODIPY staining reveal distinct LD subpopulations with either isotropic or liquid-crystalline (LC) internal organization. Scale bar, 10 µm; inset, 1 µm. **(B)** Quantification of the average LD size, number, and the fraction of polarized LDs per cell under the three conditions described in (A). Data are shown as SuperPlots, where small dots represent individual cell means, and large dots indicate the mean of each independent replicate (*N*=3). Bars display mean ± SD. Statistical significance was mainly assessed using the Mann–Whitney test, while Student’s *t*-test was employed for the Chol→OA conditions when analyzing LD size and number. For OA→Chol: WT *n*=29 cells; SKO *n*=30 cells; Chol→OA: WT *n*=37 cells; SKO *n*=33 cells; Chol&OA: WT *n*=33 cells; SKO *n*=27 cells; \*\*\**p=*0.0003.

The first notable observation was that SKO cells had only a few, but unusually large, LDs in all conditions (**Fig. 2A**), bigger than when OA was supplied exclusively. These oversized LDs showed no significant LC structures, indicating that TG accumulation was enough to dissolve LC phases by accommodating newly formed CEs (**Fig. 2B, supplementary Fig. 2A**). This finding aligns with our earlier observations and hypothesis in Figure 1, which showed that the absence of seipin promoted the dissolution of LC phases within LDs.

In WT cells, LDs were generally smaller but more numerous and exhibited higher polarization signals (**Fig. 2B, supplementary Fig. 2A**). Notably, their response varied depending on the lipid loading sequence. When OA was loaded first, or co-loaded with cholesterol, a diverse population of LC LDs appeared (**Fig. 2B, supplementary Fig. 2B**). Large hybrid LDs formed, featuring a TG-rich pocket surrounded by LC-phase CE, indicating that CE was incorporated into and assembled with LC structures, effectively displacing TG within the LD. This compartmentalization was particularly clear through the Bodipy signal, which was brighter in the TG-rich, non-LC regions of the LDs (**Fig. 2A**, *WT OA→Chol condition,* **supplementary Fig. 2B**). Additionally, some smaller LDs showed fully developed LC phases, likely due to insufficient TG incorporation during their formation, possibly due to competition with other LDs for TG recruitment. Other LDs remained in non-LC stages, highlighting the diversity of LD maturation outcomes with this loading sequence.

Initially, feeding cholesterol (+5 %OA) followed by OA, most cells displayed two well-defined LD subpopulations: on average more LDs are made, more homogeneous in size, and they are either in LC or non-LC with LC LDs tending to be smallest (**Fig. 2A-B**, *Chol→OA condition,* **supplementary Fig. 2A**). This indicates that once cholesterol initially formed LC LDs, it became more difficult, or even unfavorable, for TG molecules to incorporate into LDs afterward. TG LDs formed independently from the CE LDs, as the well-organized LC phase likely hindered TG incorporation. Overall, because TG was unlikely to efficiently integrate with LC LDs, this condition resulted in the highest number of LDs containing both LC and non-LC subpopulations (**Fig. 2B**, LD number). Consequently, LDs were the smallest across all conditions, and polarized intensity was also reduced (**Fig. 2B, supplementary Fig. 2A**).

Lastly, when we fed both OA and cholesterol simultaneously, as in Figure 1, we also observed a mixture of internal organization at equimolar loading: non-LC, half-LCs, LDs with a TG core, and the LC phase squeezed to the LD rim (**Fig. 2A**, *Chol&OA condition,* **supplementary Fig. 2B**). This condition resembles more the OA feeding conditions than cholesterol, in terms of LD number and internal organization (**Fig. 2B**).

In summary, our data suggest that, beyond the cellular TG/CE ratio, the sequence of neutral lipid buildup or the history of pre-existing LD content can improve a cell’s capacity to form new LDs or expand existing ones to store CE. As shown in Figure 1, WT cells made more LC LDs than seipin KO cells.

### The liquid-crystalline phase acts as a barrier to the incorporation of neutral lipids into LDs

Our findings show that TG and CE are fueled differently in LC and non-LC LDs. We aimed to visualize their incorporation into pre-existing LDs to better understand what drives non-LC and LC LD growth.

We used fluorescent labeling to track and measure the incorporation of newly formed neutral lipids into LDs. For TG incorporation, we employed OA labeled with 1 mol% BODIPY-tagged OA (Bpy-OA) to produce fluorescent TG (**Fig. 3A**). Similarly, for cholesterol, we added cells with cholesterol labeled with 1 mol% TopFluor-tagged cholesterol (**Fig. 3C**). The successful esterification and incorporation of the fluorescent molecules into the LDs were indicated by their integration into the LDs’ core after 24 hours of feeding (**Supplementary Fig. 3A-B**).

**Figure 3.**
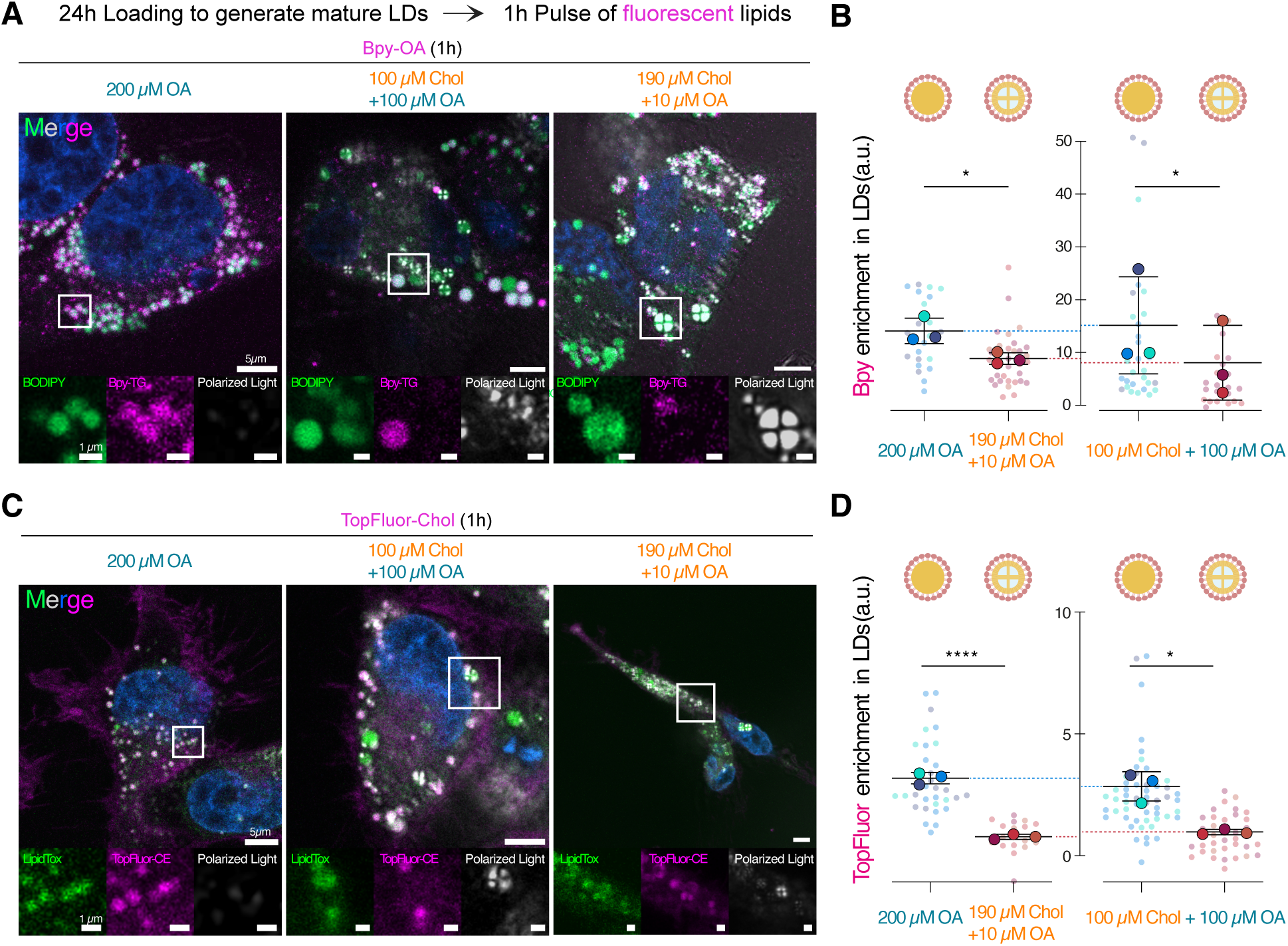
The liquid crystalline phase limits neutral lipid incorporation into lipid droplets. **(A)** Representative confocal images of HeLa WT cells preloaded for 24h with either OA or a mix of OA and cholesterol in the indicated concentrations, followed by a 1 h pulse with OA containing 1 mol% BODIPY-OA (Bpy-OA). Polarized light and BODIPY staining reveal distinct LD subpopulations with isotropic or LC internal organization. Nuclei were stained with Hoechst. Experiments were repeated three times with similar results. Scale bars, 5 µm; insets, 1 µm. **(B)** Quantification of Bpy-OA enrichment in LDs under the conditions described in (A). Enrichment of Bpy-TG was calculated as the ratio between LD and ER membrane fluorescence signals. Data are presented as SuperPlots, where small dots represent the mean value of one cell; large dots indicate the mean of each independent biological replicate (*N* = 3). LDs were separated into two groups for the equimolar condition (100 µM Chol + 100 µM OA): non-LC LDs and LC LDs. Schematic cartoons above the graphs illustrate the predominant LD organization (Isotropic vs LC). Bars show mean ± SD. Statistical significance was determined using Student’s *t*-test: 200 µM OA, *n* = 23 cells; 190 µM Chol + 10 µM OA, *n* = 35 cells, \**p* = 0.0273; 100 µM Chol + 100 µM OA non-LC LDs, *n* = 30 cells; LC LDs, *n* = 23 cells, \**p* = 0.0491. **(C)** Representative confocal images of HeLa WT cells treated as in (A) but pulsed for 1 h with cholesterol containing 1 mol% TopFluor-cholesterol (TopFluor-Chol) to monitor CE incorporation into LDs. Polarized light and LipidTox staining reveal distinct LD subpopulations with isotropic or LC internal organization. Nuclei were stained with Hoechst. Scale bars, 5 µm; insets, 1 µm. **(D)** Quantification of TopFluor-Chol enrichment in LDs under the three loading conditions. Enrichment was calculated as the ratio between LD and ER membrane signals. Data are presented as SuperPlots, where small dots represent the mean value of one cell; large dots indicate the mean of each independent biological replicate (*N* = 3). LDs were separated into two groups for the equimolar condition: non-LC LDs and LC LDs. Schematic cartoons above the graphs illustrate the predominant LD organization (Isotropic vs LC). Bars show mean ± SD. Statistical significance was determined using Student’s *t*-test: 200 µM OA, *n* = 31 cells; 190 µM Chol + 10 µM OA, *n* = 20 cells; 100 µM Chol + 100 µM OA non-LC LDs, *n* = 53 cells; LC LDs, *n* = 35 cells, \**p* = 0.0438.

We performed two separate experiments to explore the dynamics of TG and CE integration into existing LDs. First, we exposed cells to the nutrient conditions described in Fig. 1 for 24 hours, then subjected them to a 1-hour load with OA and Bpy-OA (**Fig. 3A-B**) or with cholesterol and TopFluor-Chol (**Fig. 3C-D**).

Notably, our initial observations confirmed the incorporation of both Bpy-OA and TopFluor-Chol signals into most existing LDs, although to varying extents. Under all conditions, Bpy-OA was incorporated more efficiently into LDs than TopFluor-Chol (**Fig. 3B, D**), suggesting a more straightforward incorporation of TG over CE in LDs. Further analysis revealed marked differences in the incorporation of Bpy-OA across different LD types.

Specifically, the Bpy-OA signal was more abundantly incorporated into TG-rich compared to CE-rich LC LDs, showing approximately 50% greater enrichment (**Fig. 3A-B, supplementary Fig. 3C**). This pattern persisted under equal molar loading conditions: Bpy-OA exhibited more incorporation into non-LC than LC LDs within individual cells. The enrichment levels of Bpy-OA in LC LDs under equal molar conditions closely matched those seen in CE LDs. Conversely, enrichment in non-LC LDs reflected that of TG LDs under OA loading. These results indicate that newly formed TGs tend to incorporate into isotropic LD phases, suggesting a limited flow of neutral lipids into liquid-crystalline phases.

Concurrently, we observed a markedly greater incorporation of TopFluor-Chol into TG LDs than CE LDs (**Fig. 3C-D, supplementary Fig. 3D**). The difference in TopFluor-Chol incorporation was even more striking than that observed with Bpy-OA, showing a fourfold enrichment (**Fig. 3D**). This pattern remained consistent under equimolar loading conditions. Such a pronounced disparity suggests that cells cannot effectively expand LC phases and instead preferentially channel newly synthesized CEs into TG-rich LDs. Overall, these findings corroborate our earlier observations and further support the idea that the LC phase restricts the flow of neutral lipids, impeding both their incorporation into and exit from LDs.

Lastly, we extended our neutral lipid incorporation experiments to investigate how seipin removal influences this process (**Supplementary Fig. 4A-B**). In both OA and cholesterol conditions, we found no statistically significant difference in the incorporation of Bpy-OA into existing LDs in seipin KO cells compared to WT cells (**Supplementary Fig. 4C**). This further emphasizes that the LC structure, which is less abundant in seipin KO, is a significant barrier to the incorporation of neutral lipids into LDs.

Interestingly, the lack of seipin led to a significant increase in the incorporation of TopFluor-Chol into TG LDs, with enrichment levels nearly doubling compared to WT (**Supplementary Fig. 4D**). This aligns with the observation that seipin-deficient cells led to fewer but larger LDs, which were sufficiently enriched with TG to accommodate CE. This increased TG content creates a more suitable environment for storing and dissolving newly formed CE molecules. In contrast, removing seipin did not noticeably affect TopFluor-Chol incorporation into CE LDs during cholesterol loading, where incorporation remained inherently limited (**Supplementary Fig. 4D**). All these observations were maintained under equimolar loading conditions: Bpy-OA was incorporated into LDs more efficiently than TopFluor-Chol, even in non-LC LDs (**Supplementary Fig. 4E, F**). This suggests that, at equimolar loading, non-LC LDs already contain a relatively high proportion of CE. Given this elevated baseline level, further CE addition is less favorable than TG incorporation. In other words, as the TG/CE ratio in non-LC LDs decreases, the system approaches a threshold for a transition into an LC phase. Near this threshold, additional CE incorporation becomes energetically unfavorable, whereas TG incorporation is favored, as it helps prevent entry into the LC phase.

Our findings reveal that TGs play a crucial role in incorporating CE and expanding LDs. The TG/CE ratio determines the rate and capacity of LD growth. This offers mechanistic insight into our previous observations in Figures 1 and 2. Specifically, the reduced presence of LC LDs in seipin KO cells (**Fig. 2B**) can be explained by TG-rich LDs actively drawing newly synthesized CEs from the ER membrane. As a result, the formation and growth of new LC LDs are decreased, while the growth of pre-existing TG-rich and mixed TG/CE LDs is favored.

### DGAT expression level and enzymatic activity regulate the TG/CE ratio within LD

Given TG’s specific role in regulating CE-containing LD growth, we investigated the effect of the TG synthesis enzymes on LD size and structure.

We conducted experiments at the equimolar loading condition, where the effect of TG synthesis on CE incorporation into LDs is expected to be more evident due to the high fatty acid availability. To explore the role of TG synthesis, we inhibited DGAT1, DGAT2, or both enzymes for 4 hours prior to lipid loading. We maintained the inhibition throughout the 24-hour experiment (**Fig. 4A**). When inhibiting DGAT enzymes individually, we observed a slight increase in LD size and a decrease in LD number compared to the non-inhibited condition (**Fig. 4B**). However, these differences were not statistically significant. This suggests that DGAT1 and DGAT2 may compensate for each other in producing TG to support CE incorporation into LDs.

**Figure 4.**
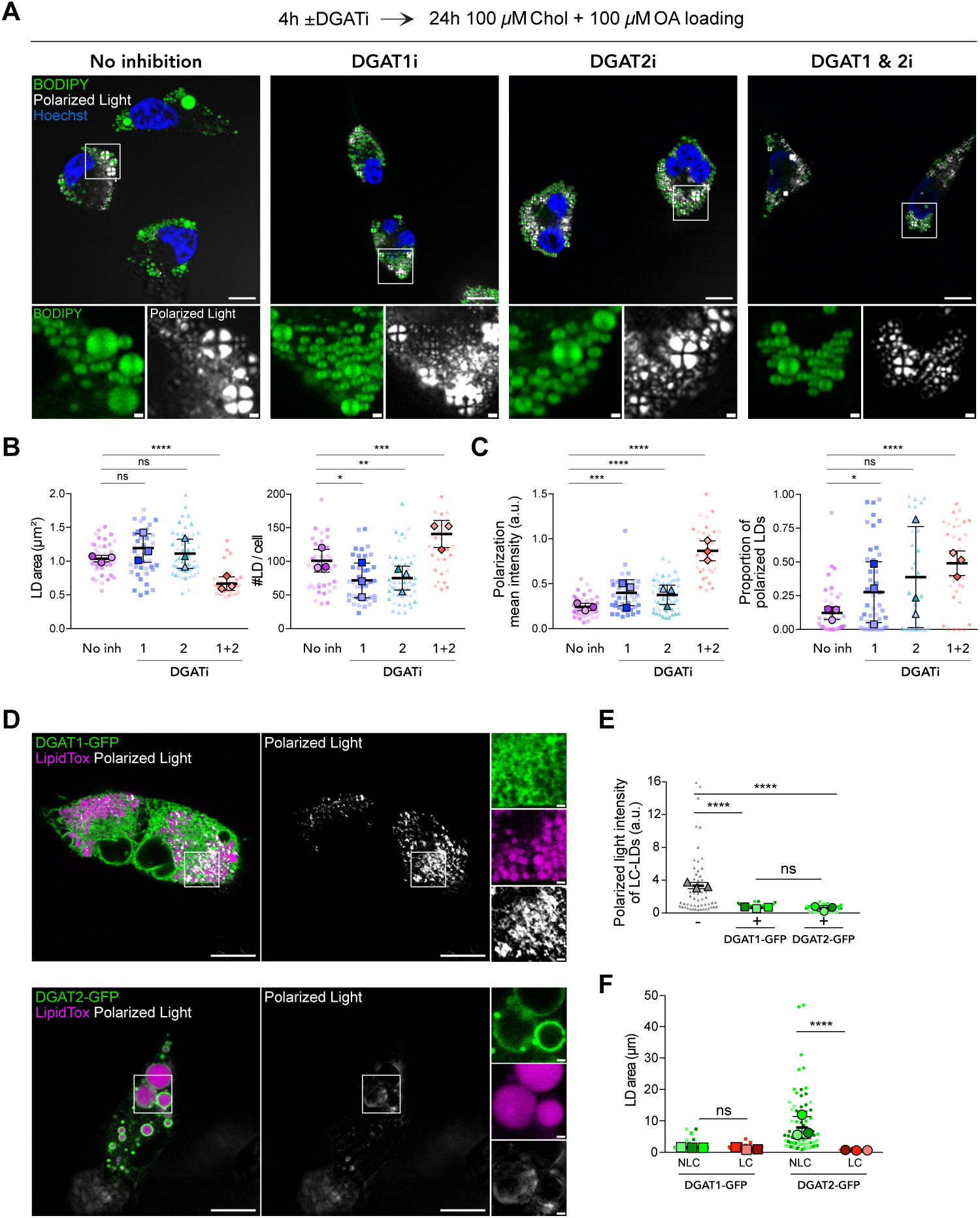
DGAT expression and activity control TG/CE ratios and lipid droplet architecture. **(A)** Representative confocal images of HeLa WT cells, either untreated (no inhibition) or treated with DGAT1 inhibitor (DGAT1i), DGAT2 inhibitor (DGAT2i), or both (DGAT1 & 2i) for 4 hours before 24 hours of equimolar lipid feeding (100 µM OA + 100 µM Chol). Polarized light and BODIPY staining reveal LD internal organization, with Hoechst staining of the nuclei. Images are representative of three independent experiments. Scale bars, 10 µm; insets, 1 µm. **(B)** Quantification of LD size and number under the indicated DGAT inhibition conditions. Each small dot represents the mean of individual cells; large dots show the mean of three independent biological replicates (*N* = 3). Bars indicate mean ± SD. Statistical significance was determined with the Mann–Whitney test: No inhibition, *n* = 39; DGAT1 i, *n* = 38; DGAT2i, *n* = 52; DGAT1 & 2i, *n* = 35. No inhibition vs DGAT1i, ns, *p* = 0. 4116; no inhibition vs DGAT2i, ns, *p* = 0. 6864; \**p* = 0. 0157; no inhibition vs DGAT2i, ** *p* = 0. 0036; no inhibition vs DGAT1 & 2i, \*\**p* = 0.0036. 0011. **(C)** Polarized mean intensity and proportion of polarized (LC) LDs under DGAT inhibition. Small dots depict individual cell measurements; large dots show the mean of three independent replicates (*N* = 3). Bars indicate mean ± SD. Statistical analysis via Mann–Whitney test: No inhibition, *n* = 40; DGAT1i, *n* = 47; DGAT2i, *n* = 35; DGAT1 & 2i, *n* = 35; ns, *p* = 0. 2105; \**p* = 0. 0245; *** *p* = 0. 0002. **(D)** Representative confocal images of HeLa WT cells transfected with DGAT1 or DGAT2, followed by 24 hours of equimolar lipid feeding (100 µM Chol + 100 µM OA). Lipid droplets (LDs) were stained with LipidTox, and internal organization was visualized with polarized light. Scale bars, 10 µm; insets, 1 µm. **(E)** Quantification of polarized light intensity in DGAT-overexpressing cells versus untransfected WT cells. Data are presented as SuperPlots: small dots show the mean of individual cells; large dots reflect the mean of three independent experiments (*N* = 3). Bars indicate mean ± SD. Mann-Whitney test results: untransfected WT, *n* = 53; DGAT1 overexpression, *n* = 162; DGAT 2 overexpression, *n* = 59; ns, *p* = 0. 4126. **(F)** LD area measurement in DGAT1- and DGAT2-overexpressing cells, categorized into non-LC (NLC) and LC LDs. Data are shown as SuperPlots, with small dots indicating the mean for a single cell and large dots showing the mean across three separate experiments (*N* = 3). Bars indicate mean ± SD. Statistical analysis with Mann–Whitney test: DGAT1 overexpression: NLC LDs, *n* = 55; LC LDs, *n* = 36; ns, *p* = 0. 3586; DGAT2 overexpression: NLC LDs, *n* = 89; LC LDs, *n* = 35.

However, the proportion of LC LDs was notably more variable across independent inhibition conditions compared to controls, and the polarized signal was, on average, more intense (**Fig. 4C**). These findings indicate that despite partial compensation, the inhibition of either DGAT isoform profoundly affects CE-rich LDs. One uncontrolled factor that may contribute to these effects is the impact of DGAT inhibition on the nucleation of CE-rich LDs^9^; thus, inhibiting either enzyme likely disrupts both the nucleation and subsequent growth of CE-containing LDs, making it difficult to disentangle their specific contributions to LD expansion.

Next, we simultaneously inhibited both DGAT1 and DGAT2. The results were striking: LC LDs became significantly more numerous, showed stronger polarization signals, and were smaller compared to LDs in either the non-inhibited or single-DGAT inhibition conditions (**Fig. 4B-C**). These findings align with our earlier observations that reduced TG synthesis promotes the formation of mature LC LDs. However, not all LDs shifted into an LC state, but this could be due to the accumulation of diacylglycerols, which may partially dissolve LC structures^9^ even under equimolar loading conditions. For comparison, under cholesterol-only loading conditions, where CE accumulation is maximized, inhibition of either DGAT individually or both together did not significantly change LD number, size, or polarization characteristics (**Supplementary Fig. 5A-B**).

Lastly, we chose to overexpress each enzyme under these equal molar feeding conditions (**Fig. 4D**). When DGAT1 was overexpressed, LC LDs still formed; however, their polarized intensity was lower compared to controls without overexpression (**Fig. 4E**). This indicates that DGAT1 activity partly reduces LC LD formation. Conversely, overexpressing DGAT2 had a much more noticeable effect: cells transfected with DGAT2 mostly showed non-LC LDs when DGAT2 localized to the LDs, and these LDs were significantly larger (**Fig. 4F**). These findings strongly support a unique role for DGAT2 in helping dissolve the LC phase, probably by encouraging direct TG synthesis and its integration into CE-enriched LDs.

Interestingly, LDs mostly exhibited LC phases in cells not transfected with DGAT2, as if DGAT2-transfected cells had specifically taken up and incorporated more OA during the same feeding period (**Supplementary Fig. 5C**). This selective enrichment was not observed with DGAT1 overexpression. The effect was even more pronounced in SKO cells (**Supplementary Fig. 5C**). Overall, our data suggest that both DGATs affect the size and internal structure of CE LDs, but DGAT2 appears to dissolve the LC phase in the LDs more effectively, likely by producing more TG locally.

### Differential binding response of perilipins to CE/TG LDs

Since apolipoproteins mark specific lipoprotein particles with distinct TG/CE ratios, thereby controlling neutral lipid transport and recruiting the right lipases, we then examined how the CE/TG ratio affects the association of the five perilipin family members. To do this, we evaluated the recruitment of each perilipin relative to its non-LD pool (**Fig. 5A-B, supplementary Fig. 6A-B**).

**Figure 5.**
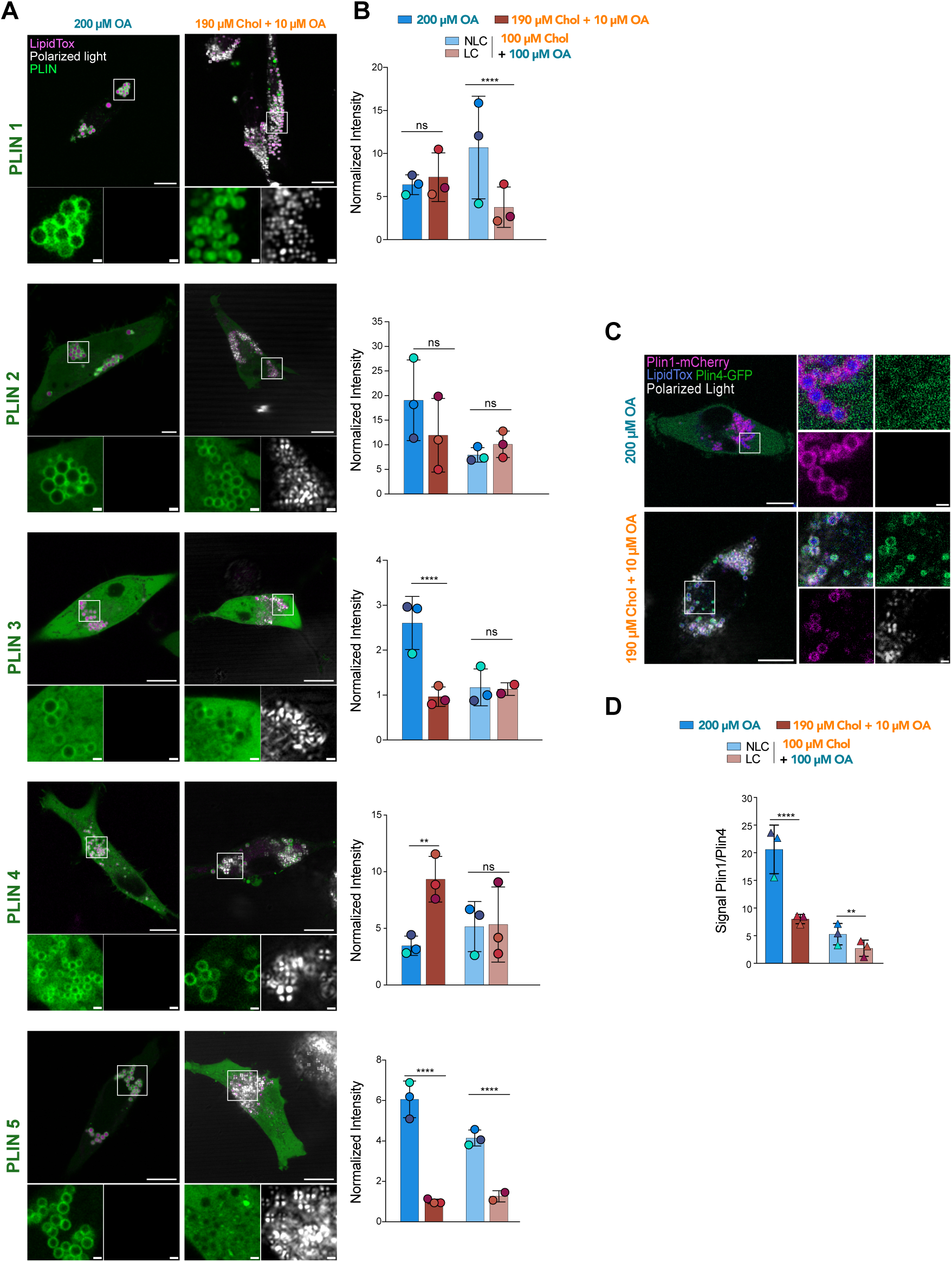
Perilipin family members exhibit distinct targeting to LDs depending on TG/CE content. **(A)** Representative confocal images of WT HeLa cells transfected with individual perilipins (Plin1– 5) for 24 hours, then loaded with 200 µM OA (TG-rich LDs) or 190 µM Chol + 10 µM OA (CE-rich LDs) for 24 hours. The internal organization of LDs was visualized using polarized light and LipidTox staining. Scale bars, 10 µm; insets, 1 µm. These images are representative of two or three independent experiments. **(B)** Quantification of perilipin enrichment on LDs under different lipid-loading conditions. In the equimolar-loading condition (100 µM Chol + 100 µM OA), two distinct LD populations were observed: LDs lacking liquid-crystalline organization (non-LC LDs, NLC) and LDs displaying liquid-crystalline phases (LC). Normalized intensity was calculated as the ratio of LD signal to non-LD pool for each cell. Each dot represents the mean of independent biological replicates (*N* = 2 or 3). TG-rich LDs: Plin1 *n*=12, Plin2 *n*=14, Plin3 *n*=21, Plin4 *n*=21, Plin5 *n*=18 cells; CE-rich LDs: Plin1 *n*=12, Plin2 *n*=14, Plin3 *n=*16, Plin4 *n*=17, Plin5 *n*=16 cells; Equimolar LDs: Plin1 *n*=14, Plin2 *n*=14, Plin3 *n*=15, Plin4 *n*=19, Plin5 *n*=24 cells. Statistical analysis was performed using the Mann—Whitney test and showed no significant differences for Plin1 (*p*=0.6450), Plin2 (*p*=0.3290 and *p*=0.2927), and Plin3 (*p*=0.1383), whereas Plin4 showed a significant difference in one condition (*p*=0.0097) but not in the other (*p*=0.9407). **(C)** Representative confocal images of co-transfected Plin1-mCherry and Plin4-GFP in cells loaded with 200 µM OA (TG-rich LDs) or 190 µM Chol + 10 µM OA (CE-rich LDs). The internal organization of LDs was visualized using polarized light and LipidTox staining. Scale bars, 5 µm; insets, 1 µm. **(D)** Quantification of co-transfection competition assays shown in (C). Normalized intensity on LDs was calculated as the ratio of LD to non-LD signal for each perilipin. Each dot represents the mean of independent replicates (*N=3*). Statistical comparisons were made using the Mann— Whitney test and highlight the dominance of Plin1 on TG-rich LDs and Plin4 on CE-rich LDs under the cholesterol feeding condition. TG-rich LDs: *n*=20; CE-rich LDs: *n*=16; Equimolar LDs: *n*=19 cells; \*\**p*=0.0067.

We observed strikingly distinct behaviors among perilipins, similar to those reported by Hsieh et al.^23^. Plin1 associated efficiently with both TG and CE LDs (**Fig. 5A-B**). Interestingly, in the equimolar TG/CE condition, Plin1 preferentially bound non-LC LDs over CE-rich LC LDs, indicating a bias toward TG storage. In contrast, Plin2 showed no such preference: it was the only perilipin that localized equally to TG and CE LDs, regardless of whether the LDs displayed an isotropic or LC phase (**Fig. 5A-B**).

Plin3 behaved differently: it bound to TG LDs but was almost entirely absent from LDs when CE was present, even under equimolar TG/CE conditions where some LDs remained isotropic (**Fig. 5A-B**). Plin4 showed the opposite trend, strongly favoring CE-rich LDs over TG-rich LDs across all feeding conditions. Finally, Plin5 exhibited the most dramatic phenotype: its LD localization was nearly completely abolished when CE accumulated, even when LDs stayed isotropic.

To further validate these differences, we performed competition assays by co-expressing pairs of perilipins. These experiments confirmed that Plin1 is the dominant TG-binding perilipin^41–43^, since it displaced Plin2-4 from TG-rich LDs (**Fig. 5C-D, Supplementary Fig. 6C-E**). Conversely, under CE-rich conditions, Plin2 could outcompete Plin1 (**Supplementary Fig. 6E**). Consistent with their CE-binding tendencies, Plin4, outcompeted by Plin1 from TG LDs, was not any more displaced by Plin1 from CE-rich LC LDs and could even displace it from the LDs (**Fig. 5C-D**).

Overall, our data support a functional specialization among perilipins: Plin1, Plin3, and Plin5 are mainly involved in TG metabolism, with Plin3 and Plin5 showing the strongest sensitivity to CE. Meanwhile, Plin4 preferentially associates with CE-rich LDs despite its ability to bind TG-rich droplets. Finally, Plin2 seems to be the regulatory perilipin unaffected by stored lipids.

## Discussion

CE and TG are the main neutral lipids in mammalian cells. A key feature of CE is its ability to form liquid crystalline phases, which are metastable liquids that tend to crystallize when contacted by a nucleator such as fibrils or cholesterol crystals^7,44–46^. Cells seem to adapt to lipid fluctuations and to regulate the innate instability of CE-rich LDs. However, this can become difficult in non-specialized cells under pathological conditions marked by excessive CE buildup, such as in hypercholesterolemia, atherosclerosis, Cholesteryl Ester Storage Disease ^47^, Wolman’s disease, Niemann-Pick disease type C, or other lysosomal storage disorders. In these cases, disrupted cholesterol metabolism and transport can destabilize CE-rich LDs, contributing to disease progression. An interesting concept in the liver is that although excessive TG accumulation is deleterious and leads to MASLD, in the context of hypercholesterolemia and cholesterol crystal formation, promoting TG biosynthesis could be protective. In this setting, diverting excess lipids toward neutral TG storage may limit cholesterol crystallization and thereby reduce cholesterol-induced cellular damage. Thus, the combination of two individually deleterious conditions may paradoxically put the brake on disease progression.

The biogenesis of TG LDs has been thoroughly studied, and the role of seipin in initiating and supporting their growth has become clearer ^16,48,49^. In contrast, much less is known about the formation and regulation of CE LDs. Recently, we identified a nucleation mechanism for CE LDs that involves a TG intermediate^9^. We found that TG buildup, especially at seipin sites^50–52^, facilitates CE LD nucleation, likely by solubilizing CE molecules, much as salt melts ice. This occurs because CE molecules, like cholesterol^53^, are prone to stack tightly^9^, losing their fluid properties. In contrast, the presence of TG promotes more favorable molecular interactions, resulting in a fluid-like material^9^. However, CE molecules within LDs eventually organize into an LC phase later as the CE/TG ratio increases. How CE LDs grow, and whether this process remains influenced by seipin and TG, remained unknown.

Here, we found that the distribution and average size of mature CE-LDs in the LC phase are not significantly impacted by seipin, indicating that the protein does not directly control their growth. This contrasts sharply with its impact on TG LDs, for which the protein removal leads to the coexistence of supersized LDs alongside numerous tiny ones, a phenomenon partly driven by TG Oswald ripening between the LDs^16,54^. In vitro, this ripening mechanism occurs only minimally in CE, which may explain why seipin has less impact on the growth of LC CE LDs. In fact, the energy needed to break the LC lattice - mainly caused by strong CE-CE interactions within the ordered onion structure inside LDs - is probably high enough to stop ripening, offsetting the internal LD Laplace pressure that is responsible for ripening^54^. Under these conditions, seipin’s counteracting role becomes less effective or unnecessary.

TG levels mainly determined the size of amorphous CE-rich LDs. When TG content exceeded around 20 mol% of the total neutral lipids^9^, the size and internal structure of LDs somewhat resembled the behavior usually seen in TG LDs. However, this effect strongly depends on the storage sequence of neutral lipids. Notably, the influence of TG was more significant when it was synthesized or added alongside or before CE accumulation. When CE LDs formed first, the later addition of newly synthesized neutral lipids - particularly CE rather than TG - became very limited, because the LC phase acted as a barrier to lipid incorporation; distinct subpopulations of CE and TG LDs develop.

The solubilization of CE by TG is a key factor that influences the size and phase state of mature CE-containing LDs, which can take different forms: pure LC phases, hybrid LC phases with a central TG pocket, or partial LC phases limited to the LD rim. Interestingly, TG’s role in destabilizing LC phases was even more pronounced in SKO cells. There, likely due to ripening, TG gradually expanded CE-rich LDs, eventually dissolving the LC structure and encouraging continuous LD growth. Thermodynamically, CE acts as a TG retention factor inside LDs because of the more favorable CE-TG interactions and the relatively unfavorable CE-CE interactions that would occur if TG were removed from the LD core^9^. This consequently increases the flux of TG toward the LD^54^. This suggests that specialized cells that handle excess cholesterol, such as steroidogenic cells, may effectively manage LDs containing CE liquid-crystalline phases. Conversely, others, such as hepatocytes, may try to prevent their formation by producing pre-existing TG-rich LDs that capture CE molecules or spontaneously triggering TG synthesis along with CE. These findings subsequently support the idea that seipin may promote the formation and stability of LC phases by fractionating TG into multiple LDs. Cells with lower seipin levels may be less inclined to form LC LDs because they can bear fewer and larger LDs containing more TGs, depending on the ER tubule/sheet ratio^18^.

Our results further underscore a critical role of TG molecules beyond their traditional function in lipid and energy storage. Other neutral lipids, such as retinyl palmitate and squalene, also require TG synthesis to efficiently incorporate into LDs^55–57^. This suggests that TG acts as a universal “LD agent,” facilitating the incorporation of diverse neutral lipids or hydrophobic molecules into LDs. Notably, the biogenesis and expansion of these LDs may not depend solely on seipin itself, but rather on the functional seipin and TG duo^9,55^.

The impact of the CE/TG ratio on LD cell biology, especially cholesterol metabolism, remains poorly understood. Studying how this ratio influences the LD proteome could provide valuable insights. Although several studies have observed changes in the LD protein composition in response to shifts in the CE/TG ratio, a detailed understanding of this remodeling process remains lacking^21^. For example, in steroidogenic cells, CE and TG may be partitioned into separate LDs, each covered by a distinct set of proteins. Our data suggest that if CE LDs form before TG synthesis, separate CE- and TG-rich LDs will coexist^23^. The TG/CE ratio is a key parameter that modulates the concentration of specific proteins associated with LDs, thereby regulating LD function.

Indeed, the TG/CE ratio characterizes lipoprotein classes and sizes, which are bound by distinct apolipoproteins that determine their lipid transport functions and lipase recruitment^28,29^. Larger particles, such as chylomicrons, have a very high TG/CE ratio and carry ApoB-48 and ApoC-II (which activates LPL). VLDLs also have a high TG/CE ratio, carrying ApoB-100 and ApoC-II to distribute endogenous TG. As VLDL is metabolized into IDL and then LDL, the TG is lost, resulting in a high CE/TG ratio for LDL, which contains only ApoB-100 for receptor-mediated cholesterol delivery. The smallest particles, HDLs, have a high CE content and contain ApoA-I (which activates LCAT to esterify cholesterol) and ApoE, both of which play key roles in reverse cholesterol transport and lipid clearance^28,29^. This indicates that apolipoproteins may have intrinsic capacities to detect and be stabilized by a range of CE/TG ratios.

Plins act as counterparts to apolipoproteins, functioning as molecular tags that coordinate LD storage and mobilization in a cell-type-specific manner. For instance, Plin1 regulates TG mobilization in adipocytes by recruiting ATGL via CGI-58 and preferentially associates with TG-rich LDs, whereas Plin2,3 must be removed from LDs to allow lipase activity^58,59^. Plin2 is unique because it is the only perilipin recruited equally to both TG- and CE-enriched LDs, suggesting a broad function in LD biology regardless of the stored neutral lipid, which may explain its widespread involvement in disorders, including infections and neurodegenerative diseases. In contrast, Plin3 and Plin5 primarily prefer TG LDs, indicating a focus on fatty acid metabolism, while Plin4 shows a preference for CE LDs, suggesting a role in cholesterol metabolism. Similar to how apolipoproteins change on circulating lipoproteins, dynamic shifts in the LD’s internal CE/TG ratio or between LDs during growth or consumption may be accompanied by a corresponding change in the surrounding Plin proteome, which ultimately defines the LD’s metabolic fate.

In conclusion, while our data illuminates the formation of mature, CE-rich LDs and confirms the essential, yet complex, role of TGs in this process, key questions remain. Future research must determine the precise moment LDs “tip over” to become CE-dominant, or LC LDs. Furthermore, we need to identify precisely which aspect of the CE/TG ratio Perilipins detect, and how the amino acid composition of each Plin promotes its specific binding preference for TG versus CE. Answering these fundamental questions is crucial to comprehending the basic metabolic and regulatory functions of LDs.

## Acknowledgment

We thank the Thiam team for the helpful discussions. French National Research Agency, ANR, ANR-24-CE11-3433-01 CHOLESTORAGE, and ANR-24-CE15-1656 MardiDROP supported this work to A.R.T. A.R.T is the recipient of the Liliane Bettencourt Prize for Life Sciences® from the Bettencourt Schueller Foundation. The Fondation pour la Recherche Médicale (FDT202504020489) and DiabetEX funded H.E.

## Author contributions

A.R.T. designed the project. H.E., C.D., and B.F. conducted all experiments and analyses with the assistance of M.O. and M.Z. C.M. and A.C. generated Plin4 plasmids. A.R.T. wrote the manuscript and reviewed it with the co-authors.

## Supplementary Figures

**Figure S1.**
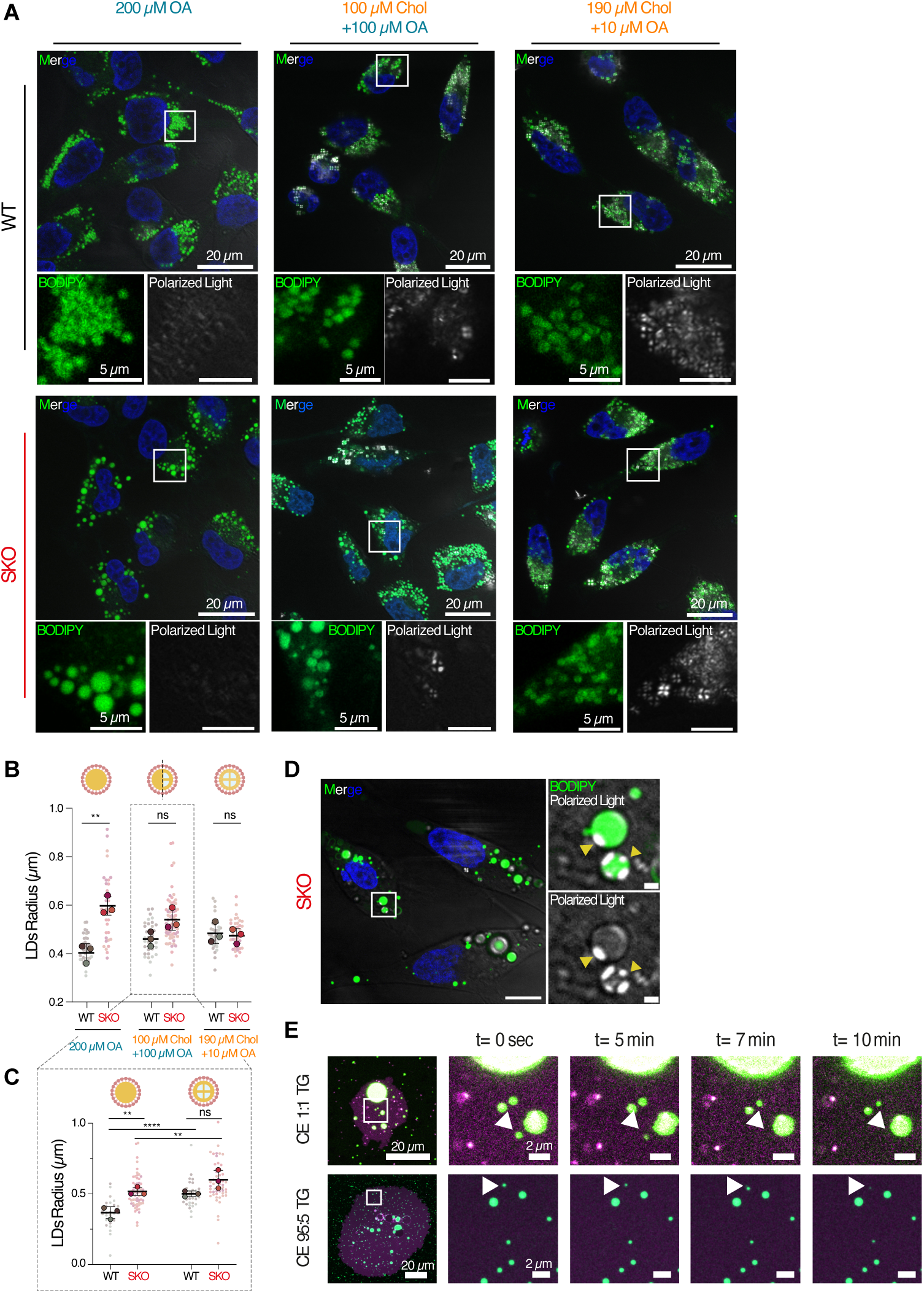
**(A)** Wide view of HeLa wild-type (WT) and Seipin knockout (SKO) cells after 24 hours of lipid loading with Oleic Acid (OA) or a combination of OA and Cholesterol at specified concentrations. This image is representative of three separate experiments. **(B)** Measurement of the average lipid droplet (LD) size per cell. Bars show the mean ± SD. Statistical analysis was done with the Mann– Whitney test. For 200 µM OA: WT *n* = 40 cells, SKO *n* = 37 cells, *p* = 0.0033; for 100 µM OA + 100 µM Chol: WT *n* = 34 cells, SKO *n* = 66 cells, ns *p* = 0.0589; for 190 µM Chol + 10 µM OA: WT *n* = 35 cells, SKO *n* = 38 cells, ns *p* = 0.7429. **(C)** Analysis of the 100 µM Chol + 100 µM OA condition, separating LDs into non-LC and LC groups. Bars represent the mean ± SD. Non-LC LDs: WT *n* = 27 cells, SKO *n* = 61 cells, *p* = 0.0068; LC LDs: WT *n* = 33 cells, SKO *n* = 51 cells, \*\**p* = 0.0015, ns *p* = 0.0649. **(D)** Confocal microscopy images of SKO cells after cholesterol treatment showing LDs with distinct internal structures. Combining polarized light microscopy with BODIPY staining reveals LD subtypes with either an isotropic pattern or a hybrid LC organization. In this example, LC LDs have a central triglyceride (TG)-rich isotropic core surrounded by a cholesterol ester (CE)-rich LC shell (zoomed area). Nuclei were stained with Hoechst. **(E)** Time course of droplet maturation in flattened vesicles containing aLDs made of equal molar TG and CE (1:1) or CE-rich mixtures (95 mol% CE / 5 mol% TG). This image is representative of three independent experiments.

**Figure S2.**
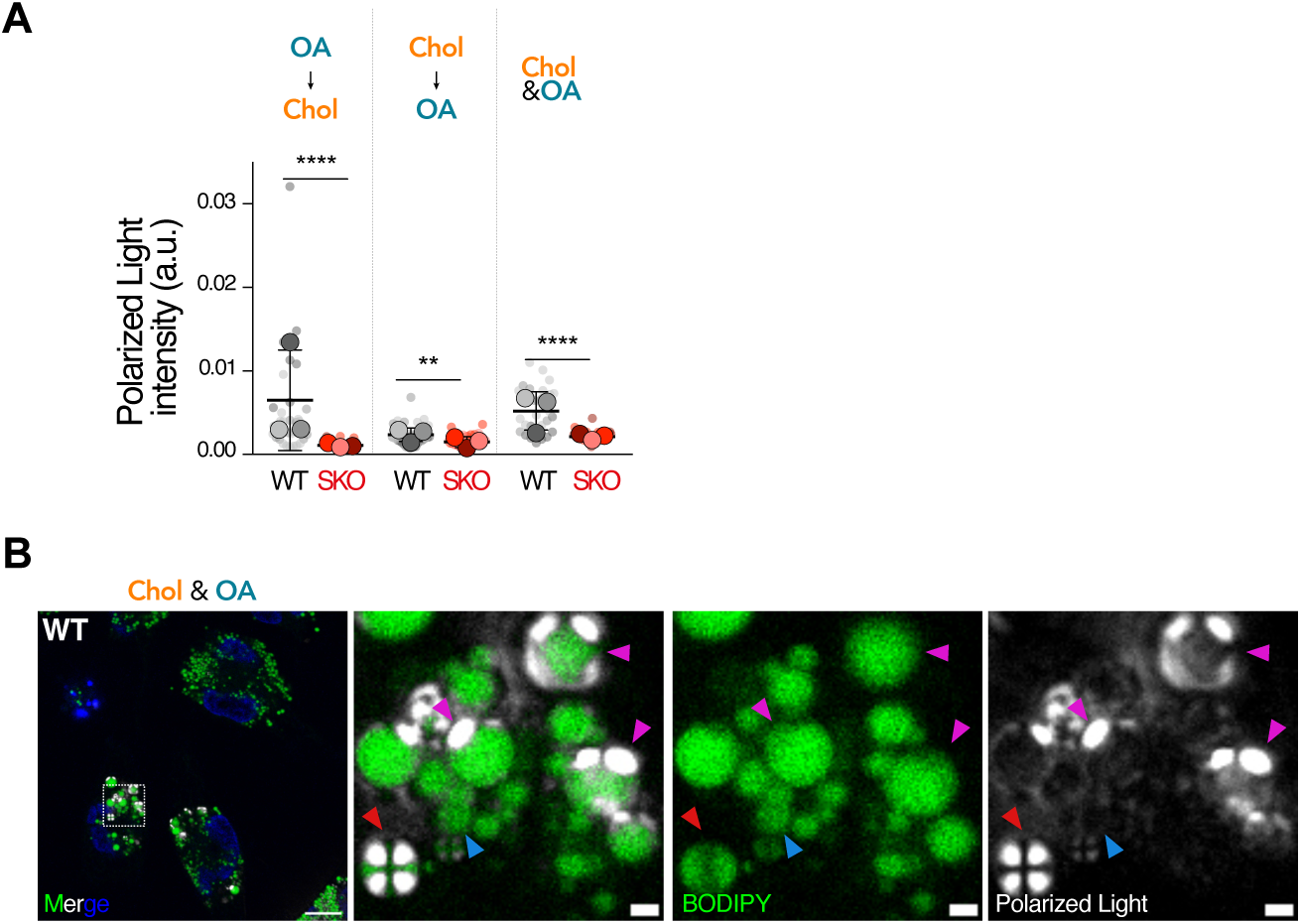
**(A)** Quantification of the LDs’ polarized light signal intensity per cell under the three conditions described in Figure 2A. Data are shown as SuperPlots, where small dots represent individual cell means, and large dots show the mean of each independent replicate (*N* = 3). Bars indicate mean ± SD. Statistical significance was tested with the Mann–Whitney test. For OA→Chol: WT *n* = 29 cells; SKO *n* = 30 cells. For Chol→OA: WT *n* = 37 cells; SKO *n* = 33 cells, \*\**p* = 0.0068. For Chol&OA: WT *n* = 33 cells; SKO *n* = 27 cells. **(B)** Confocal microscopy image of a representative HeLa WT cell shown in Figure 2A after equimolar cholesterol and OA loading, demonstrating the coexistence of multiple LD internal organizations within a single cell. Arrows point to distinct LD subpopulations: non–liquid-crystalline LDs (blue), half liquid-crystalline LDs (red), and LDs containing a TG-rich core with the LC phase compressed to the LD periphery (magenta).

**Figure S3.**
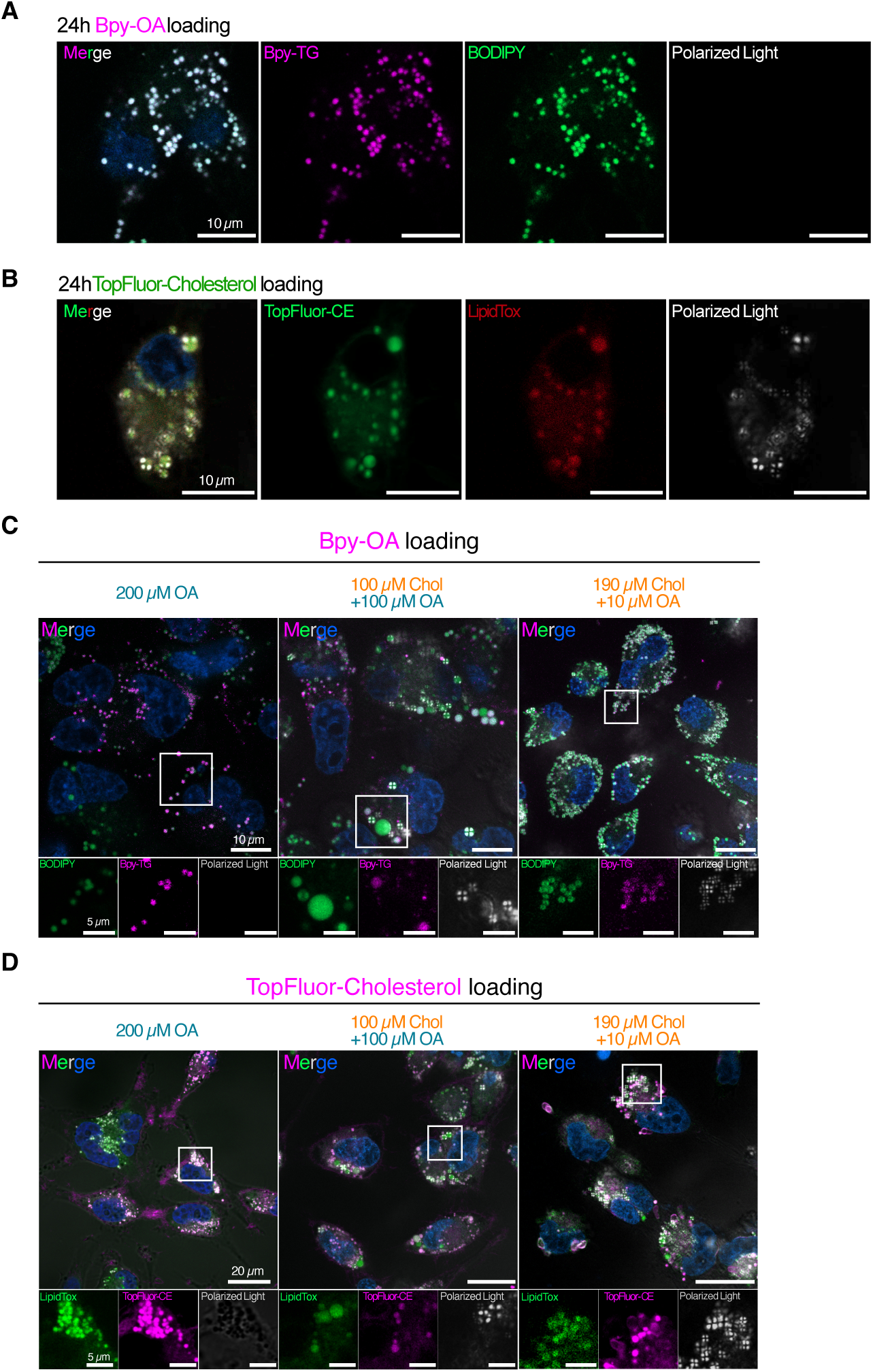
(**A-B**) HeLa WT cells were imaged after 24 hours of treatment with 200 µM Bpy-OA **(A)** or 50 µM TopFluor-Cholesterol **(B),** showing incorporation into the LDs and perfect colocalization with mature LD markers (Bodipy and LipidTox). Each experiment was repeated three times with similar results. **(C)** HeLa WT cells were imaged after 24 hours of loading with OA or a mixture of OA and cholesterol at the specified concentrations, followed by a 1-hour incubation with Bpy-OA. Experiments were repeated three times with consistent results. **(D)** HeLa WT cells were imaged after 24 hours of loading with OA or a mixture of OA and cholesterol at the indicated concentrations, followed by a 1-hour incubation with TopFluor-Cholesterol. Experiments were repeated three times with similar results.

**Figure S4.**
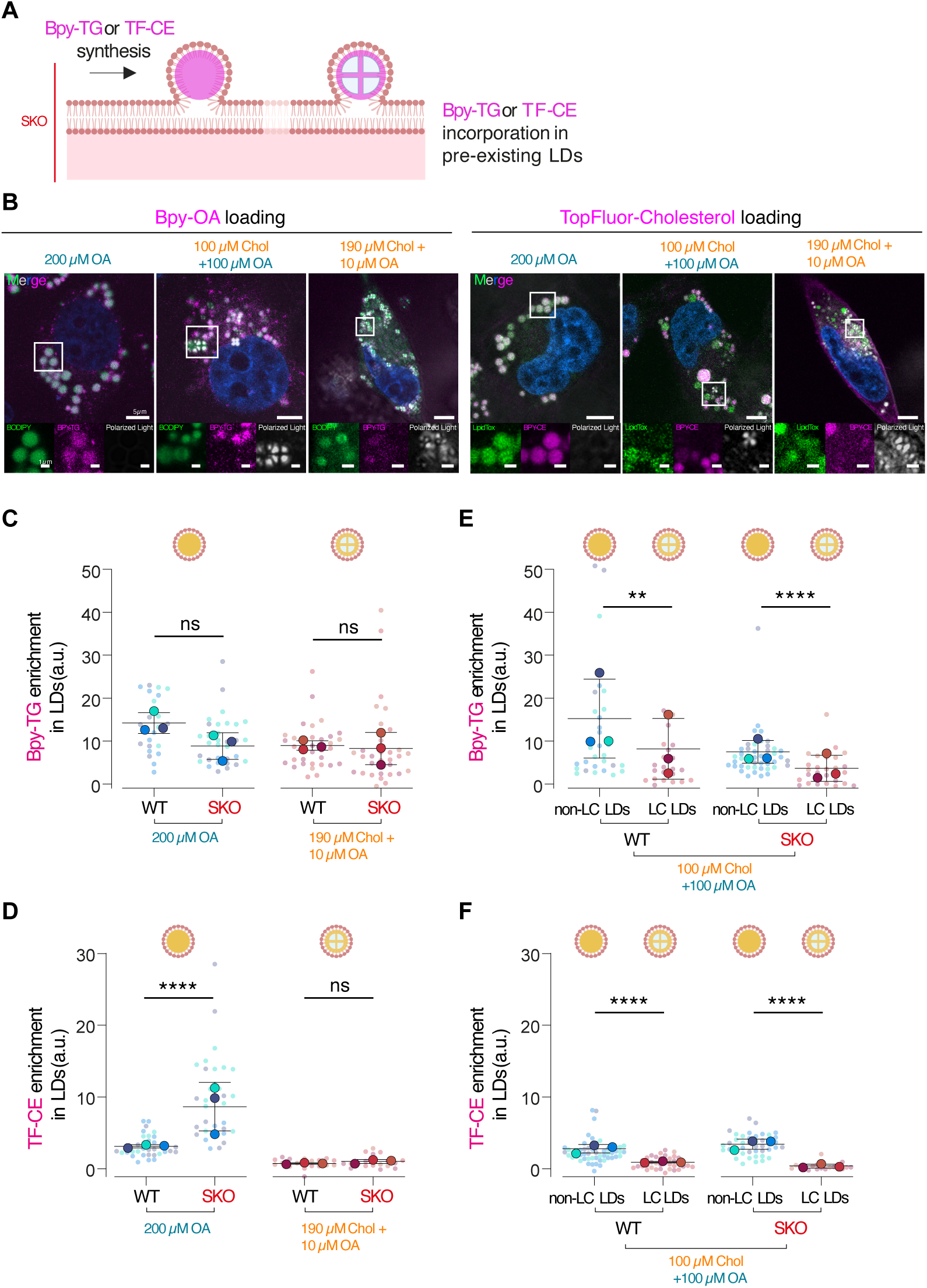
**(A)** Schematic illustration of labeled neutral lipid synthesis without seipin. SKO cells were loaded with BODIPY-OA or TopFluor-Cholesterol for 1 hour, then synthesized BODIPY-triacylglycerol (Bpy-TG) or TopFluor-Cholesteryl ester (TF-CE), respectively. The incorporation of Bpy-TG or TF-CE into existing non-LC and LC lipid droplets (LDs) was measured in the absence of seipin. **(B)** HeLa SKO cells were imaged after 24 hours of loading with OA, or a mixture of OA and cholesterol at specified concentrations, followed by a 1-hour incubation with Bpy-OA or TF-Chol, as indicated. Experiments were repeated three times with similar results. **(C)** Analysis of the conditions with 200 µM OA and 190 µM Chol + 10 µM OA from (B). Enrichment of Bpy-TG in the LDs was calculated as a ratio of the LDs’ signals to the membrane’s signals. Data are displayed as SuperPlots, with each point representing the mean from a single cell; large dots denote the mean of three independent replicates (*N* = 3). Statistical significance was tested using Student’s *t*-test for the 200 µM OA condition, and Mann–Whitney test for the 190 µM Chol + 10 µM OA condition: WT *n* = 23 cells, SKO *n* = 32 cells for 200 µM OA, with *p* = 0.0825; WT *n* = 35 cells, SKO *n* = 33 cells for 190 µM Chol + 10 µM OA, with *p* = 0.1666. **(D)** Analysis of the same conditions from (B), focusing on TF-CE enrichment in the LDs, which was calculated as a ratio of LDs to membrane signals. Each small dot marks the mean for a single cell; large dots indicate the mean of three independent replicates (*N* = 3). Statistical analysis used Student’s *t*-test or Mann–Whitney test: WT *n* = 31 cells, SKO *n* = 33 cells for 200 µM OA; WT *n* = 20, SKO *n* = 19 for 190 µM Chol + 10 µM OA, with *p* = 0.9081. **(E)** Analysis of the 100 µM Chol + 100 µM OA condition from (B). Enrichment of Bpy-TG was determined as a ratio of LD to membrane signals. LDs were categorized into non-LC and LC groups. Bars represent mean ± SD. Mann–Whitney test was used for significance: non-LC LDs, WT *n* = 30, SKO n = 45; LC LDs, WT *n* = 23, SKO *n* = 28; with p = 0.0064. **(F)** Analysis of the 100 µM OA + 100 µM Chol condition from (B), measuring TF-CE enrichment as a ratio of LD to membrane signals. LDs were divided into non-LC and LC groups. Bars show mean ± SD. Mann– Whitney test results: WT n = 53, 35; SKO *n* = 43, 18 cells for non-LC a*n*d LC LDs, respectively.

**Figure S5.**
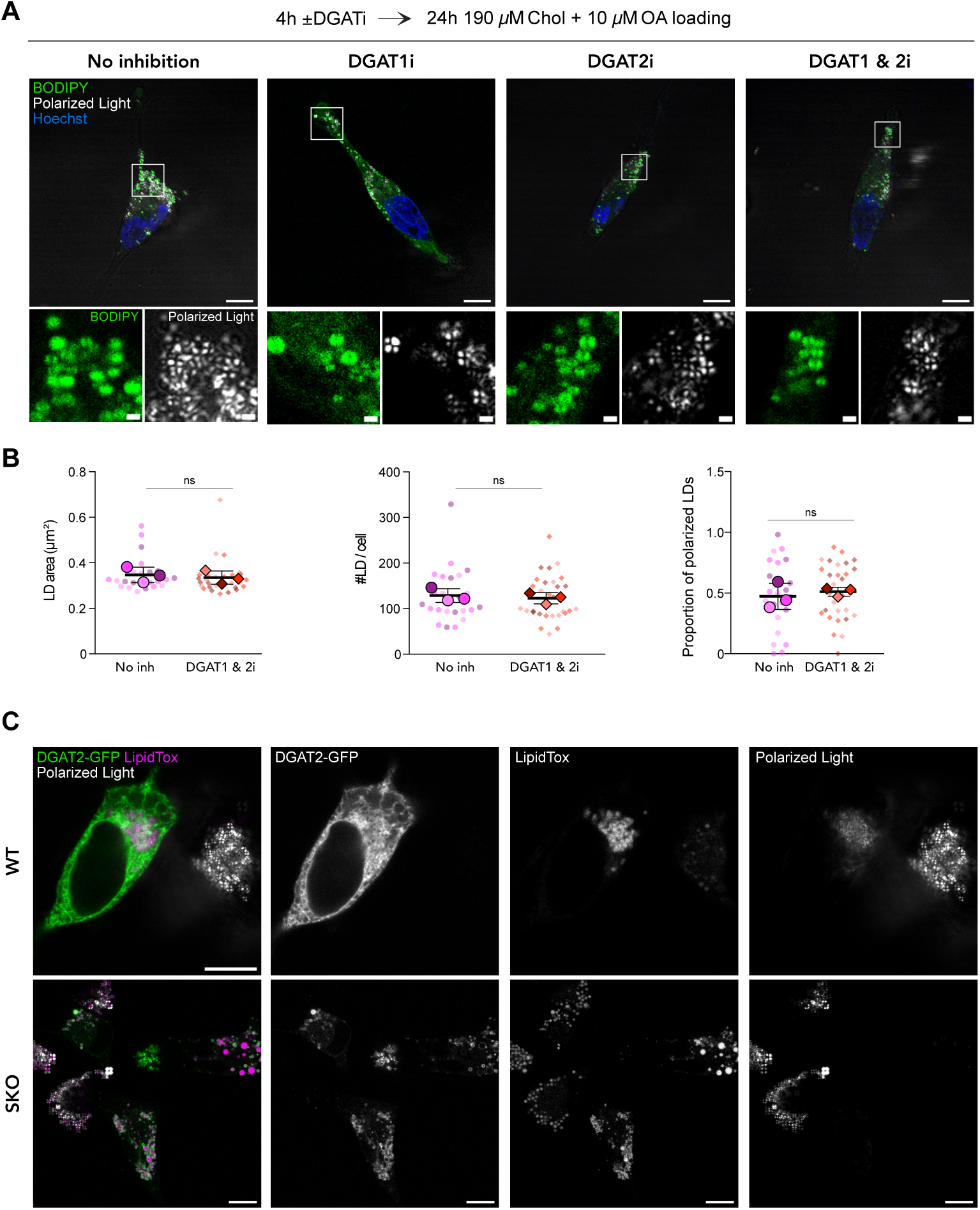
**(A)** Representative confocal images of untreated HeLa WT cells (no inhibition) or cells treated with DGAT1 inhibitor (DGAT1i), DGAT2 inhibitor (DGAT2i), or both (DGAT1 & 2i) for 4 hours before cholesterol feeding (190 µM Chol + 10 µM OA) for 24 hours. Polarized light and BODIPY staining reveal LD internal organization, with Hoechst-stained nuclei. These images represent three independent experiments. Scale bars, 10 µm; insets, 1 µm. **(B)** Quantification of LD size, number, and proportion of polarized LDs under no inhibition and DGAT1 & 2i conditions. Data are shown as SuperPlots, where each small dot indicates the mean of individual cells, and large dots show the mean of three independent biological replicates (*N* = 3). Bars represent mean ± SD. Statistical significance was assessed using the Mann–Whitney test for LD size and number, and Student’s *t*-test for the proportion of polarized LDs. No inhibition, *n* = 24 cells; DGAT1 & 2i, *n* = 30 cells. No significant differences (ns) were found: LD area *p* = 0.2639, LD number *p* = 0.6133, proportion of polarized LDs *p* = 0.6016. **(C)** Representative confocal images of WT and SKO cells transfected with DGAT2, followed by cholesterol feeding (190 µM Chol + 10 µM OA). LDs were stained with LipidTox, and their internal organization was visualized by polarized light. DGAT2-transfected cells do not display polarized LDs, in contrast to neighboring non-transfected cells within the same focal plane. Scale bars, 10 µm.

**Figure S6.**
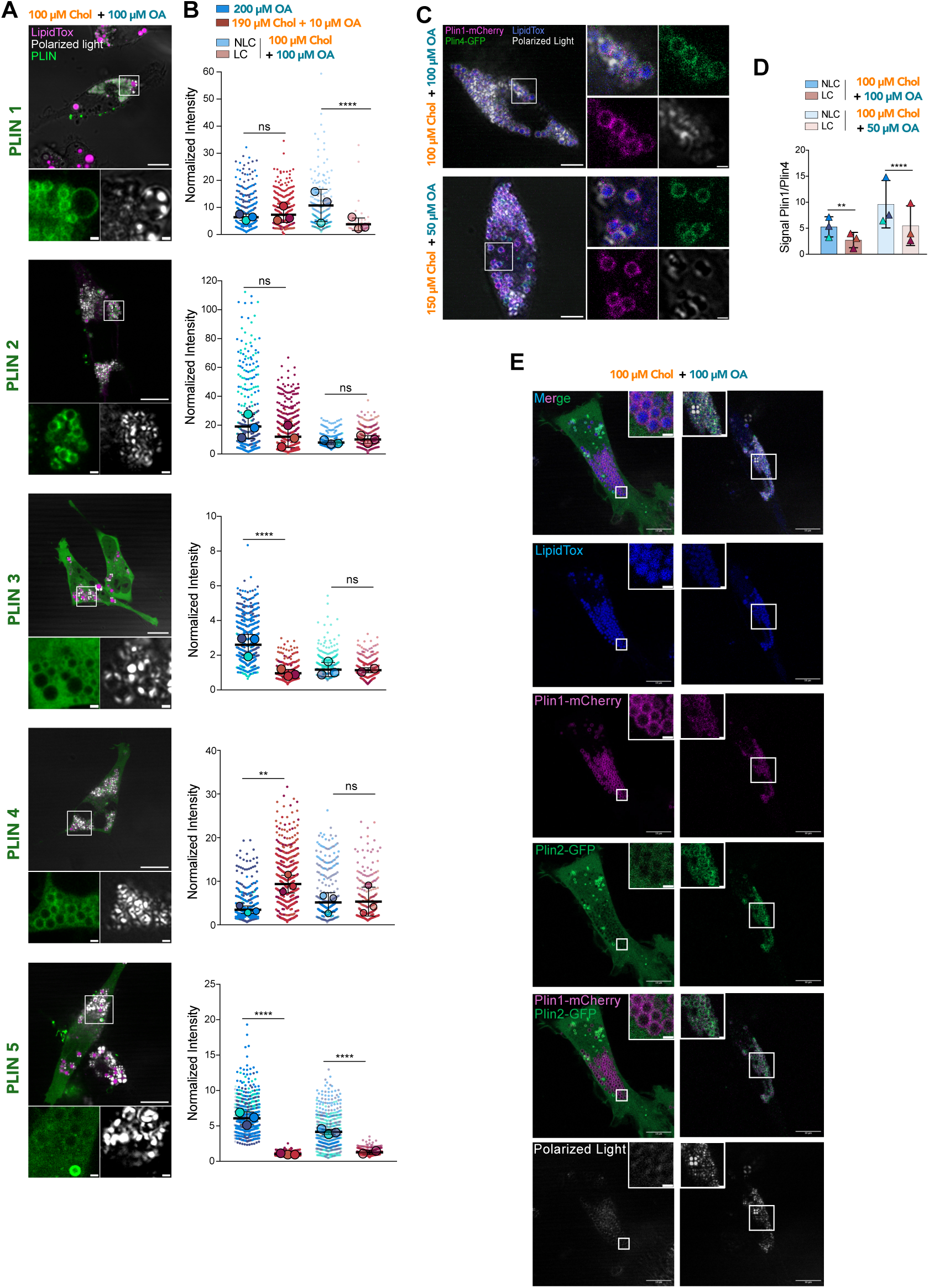
**(A)** Representative confocal images of WT HeLa cells transfected with individual perilipins (Plin1– 5) for 24 hours, then loaded with 100 µM Chol + 100 µM OA (equimolar lipid droplet, LD) for another 24 hours. The internal structure of LDs was visualized using polarized light and LipidTox staining. Scale bars, 10 µm; insets, 1 µm. **(B)** Full distribution of perilipin enrichment values across all lipid-loading conditions is shown in Fig. 5B (200 µM OA, 190 µM Chol + 10 µM OA, and 100 µM Chol + 100 µM OA). In the equimolar loading condition, two distinct LD populations were observed: LDs lacking liquid-crystalline organization (non-LC LDs, NLC) and LDs displaying liquid-crystalline phases (LC). Normalized intensity was calculated as the ratio of LD to non-LD signal for each cell. TG-rich LDs: Plin1 *n*=12, Plin2 *n*=14, ns *p*=0.2219, Plin3 *n*=21, Plin4 *n*=21, Plin5 *n*=18; CE-rich LDs: Plin1 *n*=12, Plin2 *n*=14, Plin3 *n=*16, Plin4 *n*=17, Plin5 *n*=16; Equimolar LDs: Plin1 *n*=14, Plin2 *n*=14, Plin3 *n*=15, ns p=0.1383, Plin4 *n*=19, Plin5 *n*=24. Statistical analysis was performed using the Mann—Whitney test, which showed no significant differences for Plin1 (*p*=0.6450), Plin2 (*p*=0.3290, *p*=0.2927), and Plin3 (*p*=0.1383), while Plin4 showed a significant difference in one condition (*p*=0.0097) but not in the other (*p*=0.9407). **(C)** Representative confocal images of cells co-transfected with Plin1-mCherry and Plin4-GFP loaded with 100 µM Chol + 100 µM Chol (equimolar LDs) or 150 µM Chol + 50 µM OA. The internal organization of LDs was visualized using polarized light and LipidTox staining. Scale bars, 5 µm; insets, 1 µm. **(D)** Quantification of the co-transfection competition assays shown in C. Normalized intensity on LDs was calculated as the ratio of LD to non-LD signal for each perilipin. Each dot represents the mean of three independent replicates (*N=3*). Statistical comparisons were made using the Mann—Whitney test. TG-rich LDs: *n*=20; CE-rich LDs: *n*=16; Equimolar LDs: *n*=19; 150 µM Chol + 50 µM OA: *n*=17; \*\**p*=0.0067. **(E)** Representative confocal images of competition assays between Plin1-mCherry and Plin2-GFP in cells loaded with 100 µM Chol + 100 µM OA (equimolar LDs). As in the main figure, two LD subpopulations (NLC and LC) could be distinguished by polarized light and LipidTox. Scale bars, 10 µm; insets, 1 µm.

## Materials & Methods

### Cell culture

HeLa wild-type (WT) and Seipin knockout (SKO) cell lines were maintained in high-glucose Dulbecco’s Modified Eagle Medium (DMEM, Dutscher) with 4.5 g/L glucose, stabilized glutamine, and sodium pyruvate, supplemented with 10% fetal bovine serum (FBS) and 1% penicillin-streptomycin (GibcoBRL). Cells were cultured at 37 °C in a humidified atmosphere containing 5% CO₂. For imaging, cells were seeded on 35 mm glass-bottom dishes (#P35G-0-20-C, MatTek Life Sciences) and grown for 24 h. Mycoplasma contamination was routinely checked by PCR.

### Transfection and plasmids

Transient transfections were performed using the indicated plasmids and Polyethyleneimine HCl MAX (PEI MAX, #24765, Polysciences) following the manufacturer’s instructions. Cells were transfected for 24 h prior to imaging or analysis. The plasmids used in this study include DGAT1-GFP, DGAT2-GFP by Prof. Robert Yang (UTHealth Houston, USA), PLIN1-mCherry, PLIN2-GFP, PLIN3-GFP by Prof. David B. Savage (University of Cambridge, United Kingdom), PLIN4-GFP by Alenka Čopič (CRBM, France), and Em-PLIN5 by Dr. Sarah Cohen (University of North Carolina at Chapel Hill, USA). All newly received plasmids were sequence-verified before use.

### Fluorescent probes

Nuclei were stained with Hoechst 33342 (0.025% v/v; #62249, Thermo Fisher). Lipid droplets (LDs) and membranes were labeled using HCS LipidTox™ Deep Red (0.05% v/v; #H34477) or BODIPY™ FL (0.025% v/v; #D2183).

### Chemical inhibition of DGAT enzymes

DGAT1/2 were inhibited using inhibitors (#PZ0207, #PZ0233; Sigma), dissolved in DMSO and used at a final concentration of 5 µM, for 5 h in DMEM at 37 °C under 5% CO₂.

### Preparation of loading solutions

The cholesterol/methyl-ß-cyclodextrin solution was prepared at a final concentration of 1 mM cholesterol as follows. A suitable amount of methyl-ß-cyclodextrin (#C4555, Sigma-Aldrich) was dissolved in cell culture media and then incubated with crystal cholesterol (700000 P, Avanti Polar Lipids) at a 1/8 molar ratio (cholesterol/methyl-ß-cyclodextrin) for 24 hours with agitation at 37 °C. The resulting solution was filtered through a 0.2 µm syringe filter and stored at −20 °C until use. For the preparation of labeled cholesterol (TopFluor-Chol), a mixture of TopFluor® Cholesterol (810255 P, Avanti Polar Lipids) with pure cholesterol (as described above) at a 1/100 molar ratio (TopFluor cholesterol/cholesterol) was made before complexation with methyl-ß-cyclodextrin. A stock solution (5 mg/mL) was prepared by dissolving TopFluor-Chol powder in chloroform. The chloroform was then evaporated under a desiccator, and the complexation with methyl-ß-cyclodextrin and filtration was performed as described above.

To prepare 1 mM oleic acid-containing media, mi× 10% BSA (#A8806, Sigma-Aldrich) with cell culture media at a ratio of 1/10 (v/v). Incubate the mixture with pure oleic acid (#O1383, Sigma-Aldrich) to achieve a final concentration of 1 mM. Vortex the mixture, then sonicate for 5 minutes, and incubate at 37 °C for one hour. Filter the solution using a 0.2 µm syringe filter and store at 4 °C until use. For the labeled oleic acid (Bpy-OA), prepare a mixture of BODIPY™ 558/568 C12 (#D3835, Invitrogen) with pure oleic acid at a molar ratio of 1/100 (BODIPY™ 558/568 C12/oleic acid). The remaining steps follow the same protocol as described above.

### Labeled lipids incorporation experiments

Before loading, cells’ LDs were stained either with BODIPY FL or LipidTox Deep Red. Nuclei were stained with Hoechst 33342. For labeled-OA experiments, HeLa cells’ culture medium was replaced with a solution of 200 µM of OA/BODIPY C12 in DMEM for 1 hour at the microscope. For labeled-Chol experiments, the culture medium was replaced with a solution of 50 µM of Chol/TopFluor-Chol in DMEM for 1 hour at the microscope. Lipid incorporation was visualized live at 37 °C under a confocal microscope.

### Emulsion preparation CE-containing aLDs

Cholesteryl oleate (CE, Sigma-Aldrich) was heated to 50 °C using hot water baths. 5 µL of previously liquefied CE (or a mix of CE and Triolein (TG, Sigma-Aldrich)) were vortexed for 10 seconds and sonicated for 10 seconds in 70 µL of 50 °C hot HKM buffer (containing 50 mM HEPES, 120 mM K acetate, and 1 mM MgCl2 in Milli-Q water, pH 7.4, and 275 ± 15 mOsm). It was then cooled to room temperature. For pure TG aLDs preparation, 5 µL of TG was vortexed for 10 seconds and sonicated for 10 seconds in 70 µL of room temperature HKM buffer.

### DEV preparation

Giant unilamellar vesicles (GUVs) were made of 69 mol% DOPC, 30 mol% DOPE (Avanti Polar Lipids), and 0.5 mol% Cy5-DOPE. GUVs were created by electroformation at room temperature. Phospholipids (PLs) and their mixtures in chloroform at 2.5 mM were dried onto an indium tin oxide (ITO) coated glass plate. The lipid film was dried for 1 hour. The chamber was sealed with another ITO-coated glass plate. The lipids were then rehydrated with a sucrose solution (275 ± 15 mOsm). Electroformation was performed using 100 Hz AC voltage at 1.4 Vrms for at least 2 hours. This low voltage was used to avoid hydrolysis of water and dissolution of titanium ions on the glass plate. GUVs were directly collected with a Pasteur pipette. aLDs were prepared as described above. To make droplet-embedded vesicles (DEVs), GUVs were incubated with the aLDs for 3 minutes, forming DEVs. The DEV solution was then placed on an untreated glass coverslip to induce rupture and flattening of the vesicles. BODIPY 493/503 was used to stain aLDs.

### Imaging of lipid droplets and image analysis using polarized light

Cell experiments were imaged live at either room temperature or 37 °C, using a ×60 objective on a Zeiss LSM800 microscope equipped with a polarized light visualization module. Images were initially analyzed with the segmentation “WEKA” plugin in FIJI (v2.9.0). The algorithm was trained for each experimental set and manually error-checked on test samples. WEKA produced black-and-white segmentations of the images. These images were then processed with watershed and despeckle filters before using the “Analyze particles” plugin to determine LD size and measure fluorescence levels of labeled lipids. Droplets were subsequently classified as liquid-crystalline or isotropic through cell-by-cell thresholding using a custom Python algorithm. Lipid incorporation into LDs was calculated by subtracting the fluorescent background and computing the ratio between the cell membrane fluorescent signal and droplet fluorescence.

In the manuscript, each point - transparent on graphs - is the mean fluorescence, the mean radius of the droplets, or the % of LC LDs within a cell. The mean of the points from one experiment is then used as the unique point representing the experiment and used for statistical analysis.

### Ripening analysis

Flattened DEVs were monitored for 1 hour using FIJI’s *Analyze Particles* function on the BODIPY channel. At one-minute intervals, aLDs were detected and their radii measured. The R³ value was calculated at each time point, and a linear regression was performed; the resulting slope was used as an estimate of the ripening rate.

### Data representation and statistical analysis

In most cases, SuperPlots (Lord et al., 2020) were used to show biological variability and replicate consistency, with individual cell measurements shown as small dots and the mean value per biological replicate as larger dots. Independent experiments are color-coded. Bars represent the mean of independent biological replicates ± SD, except in cases of small sample sizes, where ± SEM is displayed.

The statistical analyses were conducted using the t-test feature of GraphPad PRISM (v8.4.3), with significance assessed between two conditions via the D’Agostino and Pearson test. For normally distributed data, a two-tailed unpaired Student’s t-test was used; for non-normal data, a two-tailed Mann–Whitney test was applied. p-values less than 0.05 were considered statistically significant and are marked with asterisks (*p < 0.05; **p < 0.01; ***p < 0.001; ****p < 0.0001). Non-significant values (P ≥ 0.05) are indicated in the figure legends.

